# Chromatin remodeling of histone H3 variants underlies epigenetic inheritance of DNA methylation

**DOI:** 10.1101/2023.07.11.548598

**Authors:** Seung Cho Lee, Dexter W. Adams, Jonathan J. Ipsaro, Jonathan Cahn, Jason Lynn, Hyun-Soo Kim, Benjamin Berube, Viktoria Major, Joseph P. Calarco, Chantal LeBlanc, Sonali Bhattacharjee, Umamaheswari Ramu, Daniel Grimanelli, Yannick Jacob, Philipp Voigt, Leemor Joshua-Tor, Robert A. Martienssen

## Abstract

Epigenetic inheritance refers to the faithful replication of DNA methylation and histone modification independent of DNA sequence. Nucleosomes block access to DNA methyltransferases, unless they are remodeled by DECREASE IN DNA METHYLATION1 (DDM1^Lsh/HELLS^), a Snf2-like master regulator of epigenetic inheritance. We show that DDM1 activity results in replacement of the transcriptional histone variant H3.3 for the replicative variant H3.1 during the cell cycle. In *ddm1* mutants, DNA methylation can be restored by loss of the H3.3 chaperone HIRA, while the H3.1 chaperone CAF-1 becomes essential. The single-particle cryo-EM structure at 3.2 Å of DDM1 with a variant nucleosome reveals direct engagement at SHL2 with histone H3.3 at or near variant residues required for assembly, as well as with the deacetylated H4 tail. An N-terminal autoinhibitory domain binds H2A variants to allow remodeling, while a disulfide bond in the helicase domain is essential for activity *in vivo* and *in vitro*. We show that differential remodeling of H3 and H2A variants *in vitro* reflects preferential deposition *in vivo*. DDM1 co-localizes with H3.1 and H3.3 during the cell cycle, and with the DNA methyltransferase MET1^Dnmt1^. DDM1 localization to the chromosome is blocked by H4K16 acetylation, which accumulates at DDM1 targets in *ddm1* mutants, as does the sperm cell specific H3.3 variant MGH3 in pollen, which acts as a placeholder nucleosome in the germline and contributes to epigenetic inheritance.

## Introduction

DNA methylation, histone modification and nucleosome composition are key determinants of epigenetic inheritance, and are responsible for heterochromatin formation and transposon silencing, thereby contributing to genome stability. DDM1 was first identified in a genetic screen for loss of DNA methylation in *Arabidopsis* ^1^, and encodes a conserved SNF2-like chromatin remodeling ATPase required for both DNA and histone methylation ^2–4^. Similar reductions in both CG and non-CG DNA methylation are observed in mutants in the E3 ubiquitin ligase *VIM1 (VARIANT IN METHYLATION1*) ^5, 6^, while similar reductions in CG methylation are observed in mutants in the DNA methyltransferase *MET1* ^7^. Parallel networks have been described in mammals that utilize the DDM1 ortholog LSH/HELLS ^8–11^, the VIM1 ortholog UHRF1 (ubiquitin-like with PHD and ring finger domains 1) ^12–14^, and the MET1 ortholog DNMT1 ^15^.

Nucleosome assembly (wrapping of an H2A/H2B/H3/H4 octamer core with 1.6 gyres of dsDNA) follows DNA replication and begins with the H3/H4 tetramer, resulting in deposition of the canonical histone H3.1 by histone chaperone CAF-1 ^16–18^. This is followed by the addition of two H2A/H2B dimers ^19^. Canonical histones are subsequently replaced by histone H3.3 and histone H2A.Z at transcribed genes. This replacement requires nucleosome remodeling by SNF2 family proteins EP400 and Chd1, that alter nucleosome positioning via DNA translocation and promote both histone H3.3 and H2A.Z deposition by nucleosome unwrapping and histone exchange ^20–23^. Like other Snf2 remodelers, Chd1 binds nucleosomes at the first alpha-helix of histone H3, as well as the H4 tail, and induces a 1bp translocation per ATP cycle by distorting the DNA. The N terminus of EP400 also binds H2A.Z/H2B dimers, facilitating exchange. Specificity of Snf2 activity for genes and other chromosomal regions is conferred by recognition of histone modifications, both by the remodelers themselves as well as by subsidiary factors ^24, 25^.

Rather than promote transcription, DDM1 promotes silencing, and ATPase and nucleosome remodeling activities of DDM1 have been demonstrated *in vitro*, but only when expressed in insect cells ^26^. These activities require additional co-factors with HELLS ^27^. DDM1-dependent DNA methylation is limited to pericentromeric heterochromatin, and to transposable elements scattered along the chromosome arms. In *ddm1* mutants, these regions lose DNA methylation and associated histone H3 lysine- 9 di-methylation (H3K9me2) ^3^. Importantly, only a subset of these differentially methylated regions (DMRs) re-gain DNA methylation when DDM1 is restored, demonstrating that DDM1 is required for epigenetic inheritance ^28^. It has therefore been hypothesized that DDM1 remodels heterochromatic nucleosomes to facilitate access of DNA methyltransferases to heterochromatin ^2^. In *ddm1* mutants in Arabidopsis, and in *Lsh* mutants in the mouse, DNA methylation is lost from nucleosomes, but not from linker DNA, consistent with this idea ^29^. Arabidopsis heterochromatin comprises specific histone variants, namely H3.1, H2A.W (akin to macroH2A), and the linker histone H1 ^30^, as well as specific histone modifications including H3K27me1, H3K9me2, and deacetylated histones H3 and H4 ^31^. DDM1 impacts most, if not all, of these heterochromatin specific variants and modifications ^3, 32–35^, as does Lsh ^36, 37^, but the mechanism underlying these pleiotropic effects is unknown. We set out to determine the mechanism by which DDM1 recognizes and remodels heterochromatin, and how this contributes to epigenetic inheritance.

## Results

### DDM1 promotes the replacement of H3.3 for H3.1 in heterochromatin

Epigenetic marks in Arabidopsis are re-established during replication, when the canonical histone H3.1 is deposited and when it is specifically monomethylated on lysine-27 by ATXR5 and ATXR6 in heterochromatin ^38–40^. Over-replication of heterochromatin DNA occurs in *atxr5 atxr6* mutants, and requires *FAS2*, the CAF-1 histone chaperone ortholog that mediates H3.1 deposition ^38, 41^. Interestingly, over-replication also requires both *DDM1* and *MET1* ^42^ raising the possibility that DDM1, like FAS2, is required for deposition of H3.1. Consistently, DDM1 is expressed in dividing cells ^43, 44^ and LSH is expressed specifically during S-phase ^45^.

To test whether DDM1 plays a role in H3.1 deposition, we crossed *ddm1* mutants with H3.1-GFP and H3.3-RFP reporter lines ^39^, under the control of their endogenous promoters (HTR3 and HTR5, respectively). H3.1-GFP fluorescence was markedly diminished in root tips from *ddm1* (Figure 1A), suggesting H3.1 was depleted from chromatin. A similar loss of H3.1-GFP fluorescence was observed in root tips from *met1* (Figure 1A), as confirmed by ChIP-qPCR (Figure S1A). In *Arabidopsis*, the vast majority of H3K27me1 is specific to H3.1 and as previously reported ^46^, H3K27me1 immunofluorescence signals were also reduced in *ddm1* and *met1* to similar levels as in *fas2*^CAF1^ (Figure S1B). In contrast to H3.1, nuclei in *ddm1* had abnormal H3.3-RFP chromocenter localization suggesting ectopic deposition in heterochromatin, though the effect was less obvious in *met1* (Figure 1B; Figure S1A). To examine the role of H3 variants in epigenetic inheritance, we visualized nuclear organization of the male germline-specific H3.3 variant MGH3/HTR10, which replaces canonical H3.3 in sperm cells ^47^. MGH3 was strikingly mis-localized from the center of the nucleus to peripheral heterochromatin (defined by DAPI staining) in *ddm1* sperm cells (Figure 1C). Similar mis- localization was observed in *met1* sperm cells, but not in the non-CG methyltransferase mutant *cmt3* (Figure S1C). Importantly, mis-localization of MGH3 in sperm cells was observed even when DDM1 or MET1 function was restored in heterozygotes (86.7%±5.77 of pollen grains from MGH3-GFP X *ddm1* F1 plants; 96.7%±5.77 from MGH3-GFP x *met1* F1 plants; n=3 heterozygous plants each). This indicated that MGH3 mis-localization was somehow inherited from *ddm1* mutants, even when DDM1 function was restored.

**Figure 1.**
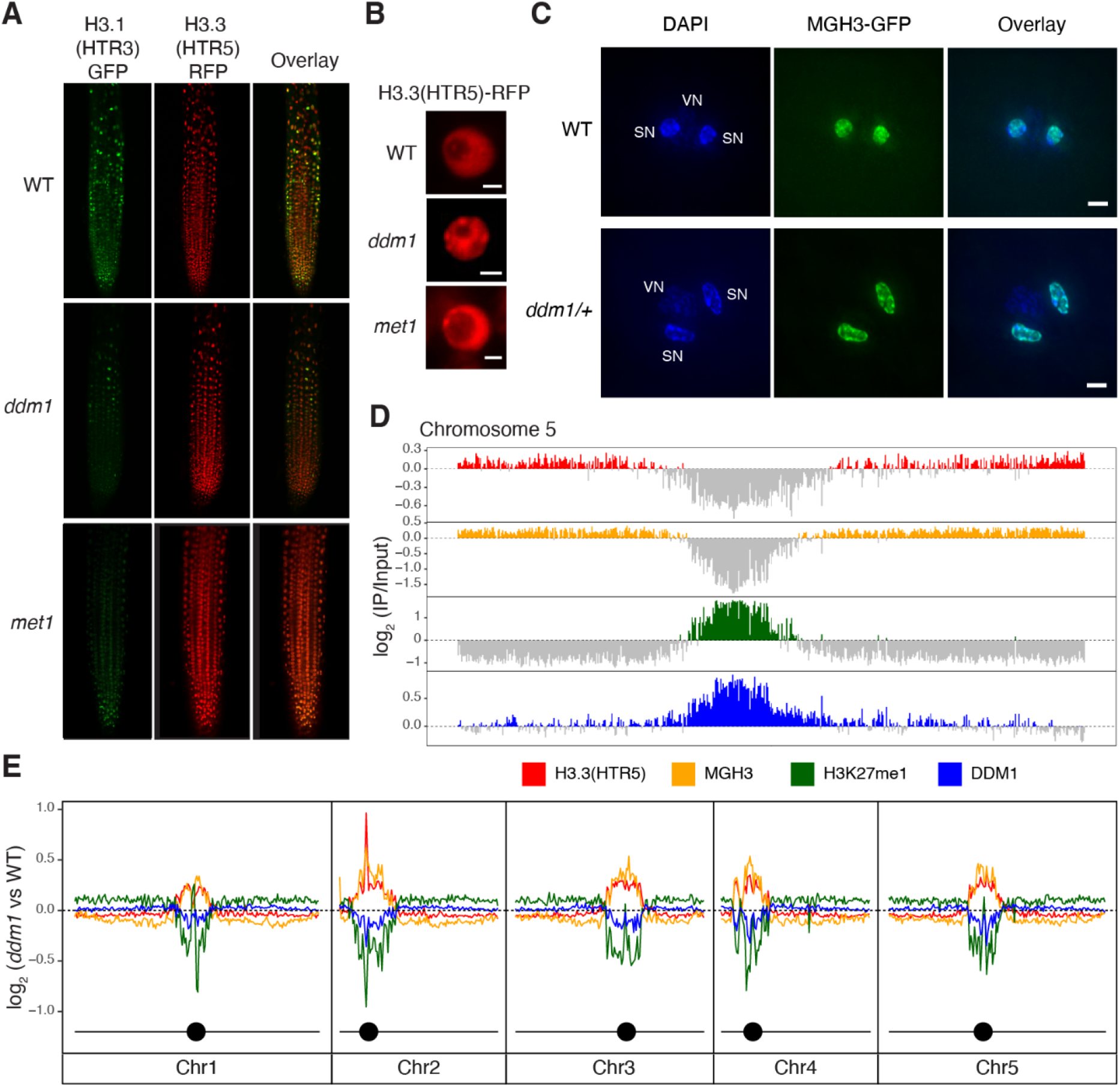
Replacement of histone H3.1 by H3.3 in *ddm1* mutants. (A) H3.1(HTR3)-GFP and H3.3(HTR5)-RFP localization in Arabidopsis root tips of wild-type (WT), *ddm1*, and *met1*. (B) Ectopic chromocenter localization of H3.3(HTR5)-RFP in *ddm1* as compared to WT and *met1*. Scale bars indicate 2 μm. (C) Male <u>G</u>ermline-specific <u>H</u>istone H3.3 variant MGH3-GFP localization in sperm nuclei of Arabidopsis pollen grains from WT and *ddm1*/+ plants. DAPI staining was used to visualize vegetative (VN) and sperm nuclei (SN). Mis-localization to the nuclear periphery was observed in pollen from *ddm1*/+. Scale bars indicate 2 μm. (D) Distribution of ChIP-seq marks in WT along chromosome 5, showing preferential localization of H3.3(HTR5) in leaf tissue and MGH3 in pollen on chromosome arms, and H3K27me1 and DDM1 on pericentromeric heterochromatin. The values correspond to the log2 fold change of IP/H3 for H3.3(HTR5) and H3K27me1, and IP/Input for DDM1 and MGH3, normalized in counts per million. Signal tracks were averaged in 50kb windows with negative log2 values shown in grey. (E) Distribution of the log2 ratio of the ChIP-seq coverage between *ddm1* and WT, showing an increase in H3.3 and MGH3 in peri-centromeric regions, coupled with a loss of H3K27me1 and DDM1. MGH3 IP was performed on pollen grains from a heterozygote *ddm1*/+ plant (as in C). In *ddm1-2* mutants, DDM1 protein is present at reduced levels (Figure S1E).

Many of the phenotypes observed in *ddm1* mutants are also found in *met1*, including the epigenetic inheritance of transposon DNA hypomethylation at CG dinucleotides ^48^. There are some important differences however, such as the loss of CHG methylation from transposons in *ddm1*, and the loss of gene body CG methylation from exons in *met1* ^48^. One explanation could be that DDM1 and MET1 interact and depend on each other for accumulation in heterochromatin. To further explore interactions between DDM1 and MET1 we expressed a functional MET1-mCherry fusion in transgenic plants (see Methods) that also expressed a functional DDM1-GFP fusion ^44^ and observed co- localization in interphase (Figure S1D). We then performed bimolecular fluorescence complementation by transient expression of split GFP fusion proteins in Arabidopsis leaves, and detected robust complementation indicating close proximity in the nucleus (Figure S1E). Finally, we raised polyclonal antibodies to DDM1 (see Methods) and determined by Western blotting that levels of DDM1 protein were sharply reduced in *ddm1* mutants (Figure S1F), consistent with a splice donor site mutation in *ddm1-2* ^49^. Importantly, *met1-1* mutants had a similarly low level of DDM1 protein in chromatin fractions compared to wild-type (Figure S1F). In mammalian cells, Dnmt1, Lsh and Uhrf1 interact, and are co-recruited to the replication fork by a combination of histone modifications and hemi-methylated DNA ^10, 50–53^. Our results support a similar interaction between Arabidopsis orthologs MET1, DDM1 and VIM1. We therefore focused our subsequent studies on DDM1.

First, we investigated the genome-wide effects of DDM1 on nucleosome composition and modification by ChIP-seq, using antibodies against H3K27me1 and antibodies against H3.3(HTR5)-RFP, as well as by low-input Chip-seq of MGH3-GFP from pollen (Figure 1D). As expected, H3K27me1 was confined to pericentromeric heterochromatin, while H3.3 and MGH3 were found in the gene dense chromosome arms ^40, 54^. We also performed ChIP-seq using anti-DDM1 antibodies (see Methods), and found that DDM1 was found in pericentromeric regions, overlapping closely with H3K27me1, precisely where H3.3 was depleted (Figure 1D; Figure S1G). Next, we examined *ddm1* mutants by ChIP-seq using the same antibodies. Consistent with microscopy-based observations, H3K27me1 was depleted in *ddm1* compared to wild-type (Figure 1E), while H3.3 was ectopically deposited in pericentromeric regions (Figure 1E). Low-input Chip-seq of pollen from MGH3-GFP X *ddm1* F1 plants using anti-GFP antibodies revealed that MGH3 was also ectopically deposited in heterochromatin in pollen (Figure 1E), consistent with peripheral nuclear localization in sperm cells (Figure 1C).

### H3.3 deposition prevents DNA methylation of heterochromatin in *ddm1* mutants

It has previously been shown that DDM1 is required for methylation of nucleosomal DNA, but not for linker DNA, suggesting that DDM1 allows access to DNA methyltransferase by remodeling the nucleosome ^29^. The loss of H3.1 and gain of H3.3 in *ddm1* mutants (Figure 1E) suggested that H3.3 might prevent methylation when DDM1 was removed. We therefore investigated whether loss of histone H3 variants or their chaperones could rescue loss of DNA methylation in *ddm1* mutants. *fas2* encodes the large subunit of the CAF-1 histone chaperone responsible for H3.1 deposition during S phase. We did not obtain *ddm1 fas2^CAF^*^1^ double mutants (n=150 F2 plants), while the siliques of *ddm1*/+ *fas2^CAF^*^1^ plants contained undeveloped seeds (Figure 2A; Methods). This can be explained by exacerbated loss of H3.1 in *ddm1 fas2^CAF^*^1^ double mutants resulting in synthetic lethality.

**Figure 2.**
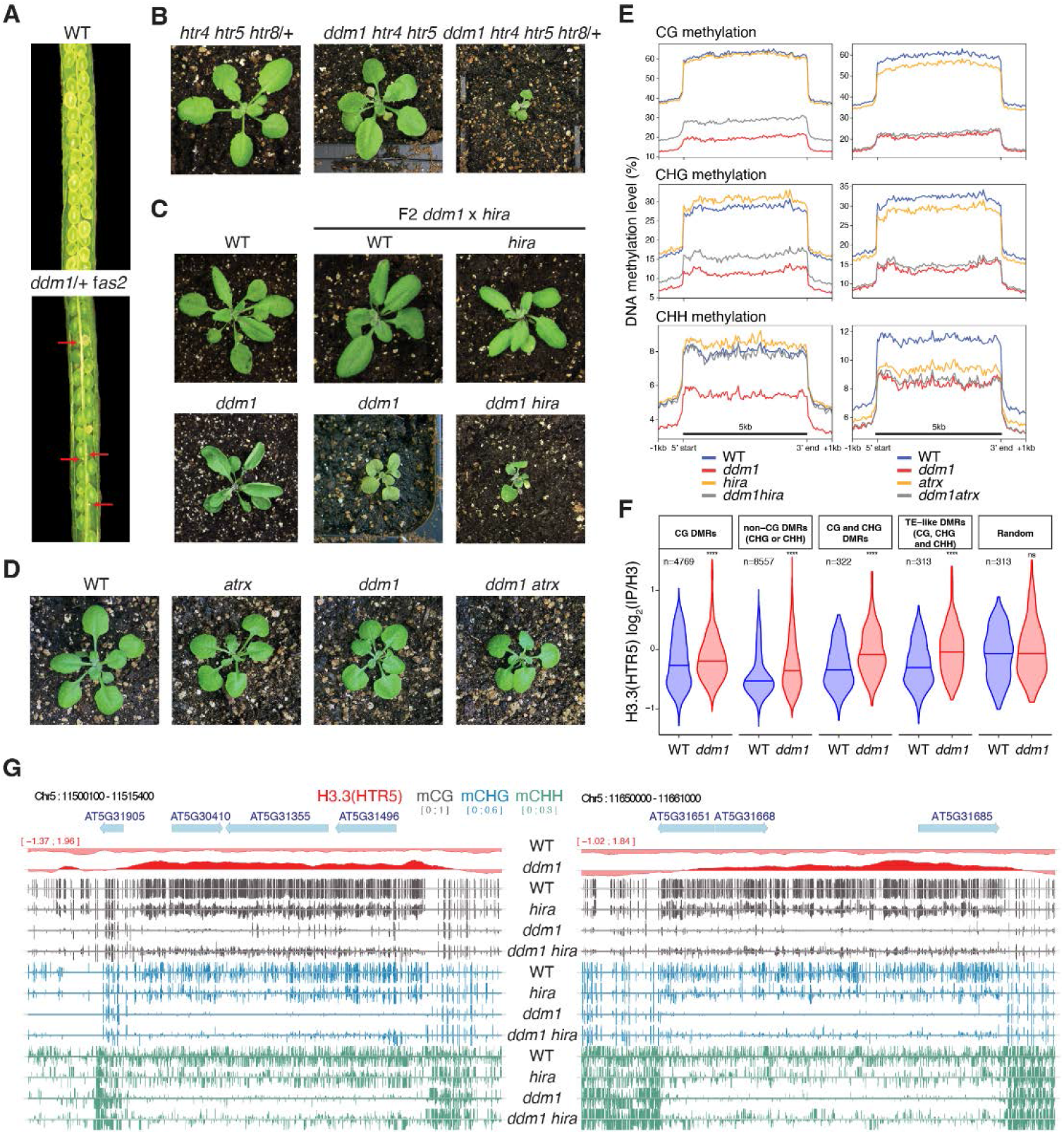
Genetic interactions between *ddm1*, histone H3 variants and chaperones impact DNA methylation. *fas2* and *hira* are mutants in H3.1 (CAF-1), and H3.3 (HIRA) chaperones, respectively. ATRX is a chromatin remodeler required for H3.3 deposition. (A) Siliques of wild-type (WT) and *ddm1*/+ *fas2* plants. Red arrows indicate nonviable seeds (synthetic lethality). (B) F2 *ddm1 htr4 htr5 htr8*/+ with reduced H3.3 has severe growth phenotypes compared to *htr4 htr5 htr8*/+. (C) F2 *ddm1 hira* double mutants from *ddm1* and *hira* parents, compared with WT, *hira* and *ddm1* siblings. (D) F2 *ddm1 atrx* double mutants from *ddm1* and *atrx* parents compared with WT and *ddm1* siblings. *ddm1 hira* and *ddm1 atrx* were phenotypically indistinguishable from *ddm1* siblings but *ddm1 hira* were more severe. (E) DNA methylation levels in CG, CHG and CHH contexts in *ddm1* and *hira* mutants on the left, and *ddm1* and *atrx* mutants on the right, determined by whole genome bisulfite sequencing. The DNA methylation levels range from 0 to 100% and are substantially increased in *ddm1 hira* as compared to *ddm1*. *atrx* mutants lose some methylation and fail to rescue methylation loss in *ddm1*. Metaplots calculated from all 31,189 transposable elements annotated in TAIR10. (F) Levels of H3.3 in WT and *ddm1* Chip-seq at differentially methylated regions (DMRs) between *ddm1* and *ddm1 hira* (hyper-methylated in *ddm1 hira*). The number of DMRs (n) in the different cytosine nucleotide contexts are noted. H3.3 is statistically enriched in *ddm1* compared to WT at these DMRs, but not in random regions (**** P<0.0001, ns not significant, t-test). See Table S1 for the list of all DMRs. (G) Representative loci that re-gain DNA methylation in *ddm1 hira* as compared to *ddm1*. Ectopic H3.3 in *ddm1* is shown above (red track).

To assess the role of H3.3 in DNA methylation, we made double mutants between H3.3 and *ddm1*. The complete knock-out of all H3.3 genes (*htr4, htr5,* and *htr8*) is lethal in Arabidopsis ^55^ and we obtained no viable *ddm1 htr4 htr5 htr8* mutants. *ddm1 htr4 htr5 htr8*/+ mutants, which had only one functional copy of H3.3, were slow growing while *ddm1 htr4 htr5* mutants, which had 2 functional copies, were normal indicating dose dependence (Figure 2B). As an alternative to using H3.3 mutants, we examined interactions with mutants in the H3.3 chaperones HIRA and ATRX. The transcriptional histone chaperone HIRA is required for H3.3 deposition during interphase, and has viable mutants ^17, 18, 56, 57^. *ddm1 hira* double mutants exhibited delayed growth phenotypes, but these were comparable to *ddm1* siblings and were also viable (Figure 2C). ATRX encodes a conserved Snf2-like remodeler specifically required for H3.3 deposition in heterochromatin just before mitosis ^58–62^. By contrast with *ddm1*, double mutants between *atrx* and *hira* are inviable while *atrx fas2^CAF^*^1^ double mutants are viable ^62^ and we found that *atrx ddm1* double mutants were also fully viable (Figure 2D). These contrasting phenotypes with H3.1 and H3.3 chaperone mutants are consistent with DDM1/FAS2 and ATRX/HIRA having essential roles in H3.1 and H3.3 deposition, respectively.

Next, we performed whole genome bisulfite genome sequencing of *ddm1, atrx* and *hira* mutants. DNA methylation levels at transposable element loci were dramatically reduced in *ddm1* siblings for all methylation contexts, as expected, but methylation levels were significantly higher in *ddm1 hira* double mutants (Figure 2E). In contrast, *ddm1 atrx* mutants did not recover DNA methylation, while *atrx* single mutants actually lost methylation (Figure 2E). In order to determine if H3.3 was responsible, we examined H3.3 levels at differentially methylated regions (DMRs) that recovered DNA methylation in *ddm1 hira*, and found a highly significant enrichment of H3.3 in *ddm1* mutants relative to wild-type (P<10e-14; Figure 2F). Significant enrichment for H3.3 was found at DMRs genome wide (Figure 1E; Figure 2F) and at individual loci, some of which completely lost DNA methylation in *ddm1* mutants but recovered substantially in *ddm1hira* (Figure 2G). These results indicated that ectopic deposition of H3.3 was responsible for loss of methylation in *ddm1* mutants, while ectopic H3.1 deposition was presumably responsible for loss of methylation in *atrx* mutants. In both cases, loss of DNA methylation occurs because nucleosomes block access to DNA methyltransferase in the absence of chromatin remodeling ^29^.

### Single-particle Cryo-EM reconstruction of the DDM1-nucleosome complex reveals interactions with DNA and with histones H3 and H4

To establish the molecular specificity of physical interactions between DDM1 and nucleosomes, the molecular structure of a DDM1-nucleosome complex was determined by single-particle cryo-electron microscopy (cryo-EM). A nucleosome core particle comprised of H2A.W, H2B, H3.3, H4, and 147 bp Widom 601 DNA was assembled with full-length DDM1 (Figure 3A). After iterative rounds of filtering, classification, and refining selected particle classes (Figure S2A), a 3D reconstruction of the DDM1-nucleosome complex was obtained at 3.2 Å resolution (Figure 3B; Figure S2B-D). Estimation of the local resolution was higher for the nucleosome core compared to DDM1 (Figure S2C,D). The final structure spans residues 200-435 of DDM1, including the DEXD ATPase domain, and residues 442-673, which include the helicase superfamily C-terminal (HELICc) domain (Figure 3A).

**Figure 3.**
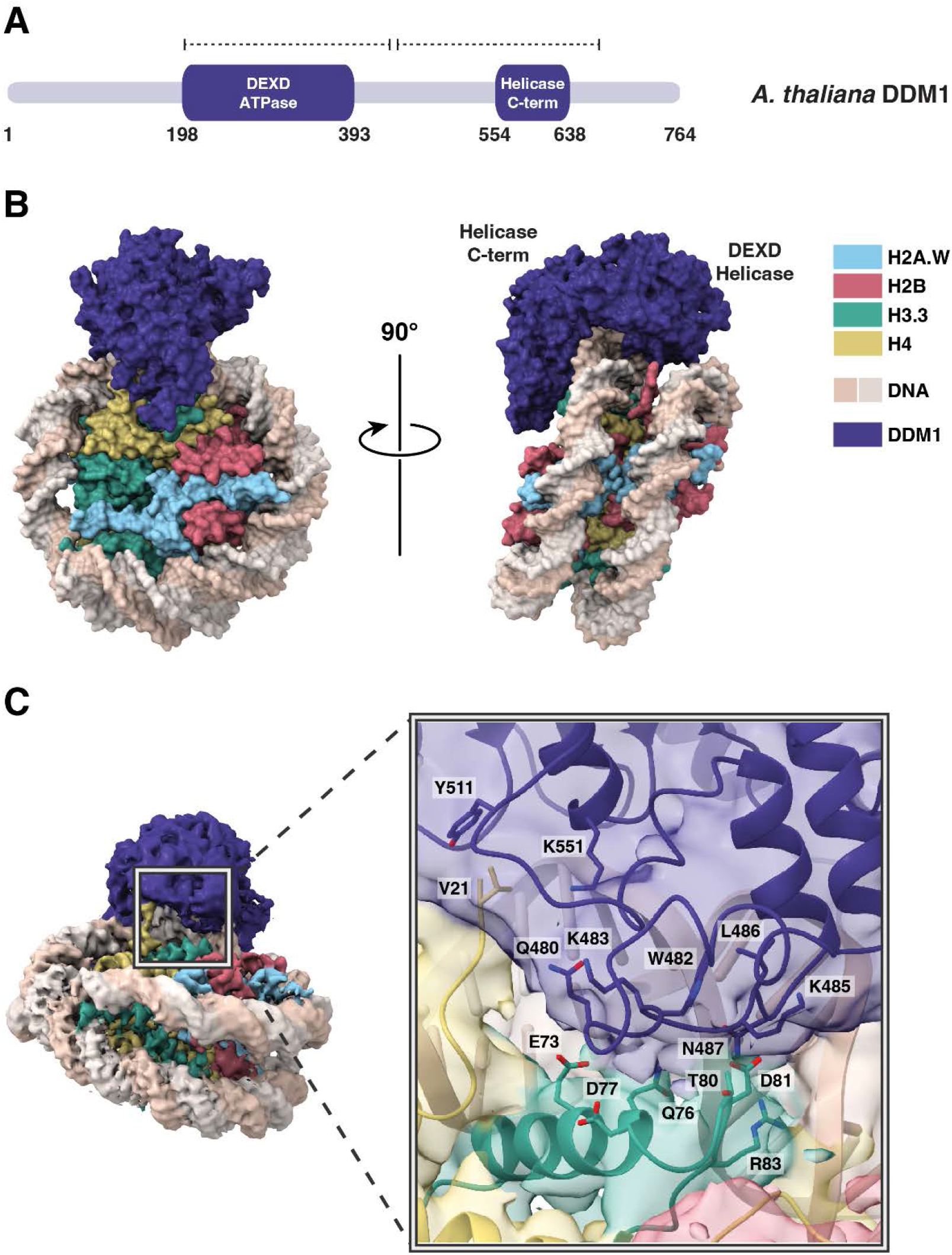
Structural basis of DDM1-nucleosome interactions. (A) Protein domain schematic of DDM1. Residue numbers indicate the boundaries of DDM1 and its domains: the N-terminal DEXD ATPase domain and helicase superfamily C-terminal domain (HELICc). Dashed lines represent the coverage of the DDM1 molecular model. (B) Overview of the molecular structure of the DDM1-nucleosome complex as determined by cryo-EM. DDM1 domains, corresponding to the two lobes, are labeled on the side view. (C) DDM1-histone interactions. The experimental density map of the complex shows that DDM1 interacts with histones H3.3 (green) and H4 (yellow). For the inset, a cartoon representation with partially transparent cryoEM map colored by domains is shown. Amino acids along the DDM1-histone interface (6 Å cutoff) are displayed as sticks, and include T80 and D81 of histone H3.

Like other Snf2 remodelers, DDM1 has two main lobes, each consisting of a parallel β- sheet core surrounded by short a helices, connected by a flexible linker (Figure 3B; Figure S3A,B). As expected from intrinsic disorder predictions, the N-terminal DEXD ATPase domain and the C-terminal HELICc helicase domain exhibited higher resolution than the peripheral regions (Figure S2D). Density for the disordered N-terminus was not observed in the cryo-EM map (Figure 3A), consistent with predictions of a highly disordered domain.

DDM1 clasps the nucleosome on the outside of superhelix location-2 (SHL-2), making contact with both gyres as well as the histone octamer core. By comparison with other Snf2-family structures ^63–66^ this placement of DDM1 and the associated distortion of the DNA helix indicates a role in DNA translocation, nucleosome sliding and assembly or disassembly (Figure S3A-C). Both lobes of DDM1 make multiple contacts with the DNA, with positively charged grooves and patches of DDM1 serving as DNA interfaces (Figure S3D). The helical structure of DNA was notably unwound where the upper gyre contacts the HELICc domain, and the DNA strands displaced about 7 Å toward the enzyme, while the other gyre shifts in the other direction compared to an unbound nucleosome, causing an opening between the two gyres (Figure 4A). This feature is observed in other chromatin remodelers ^67, 68^, in both nucleotide-free and ADP-bound states, though the displacement for DDM1 appears to be somewhat larger.

**Figure 4.**
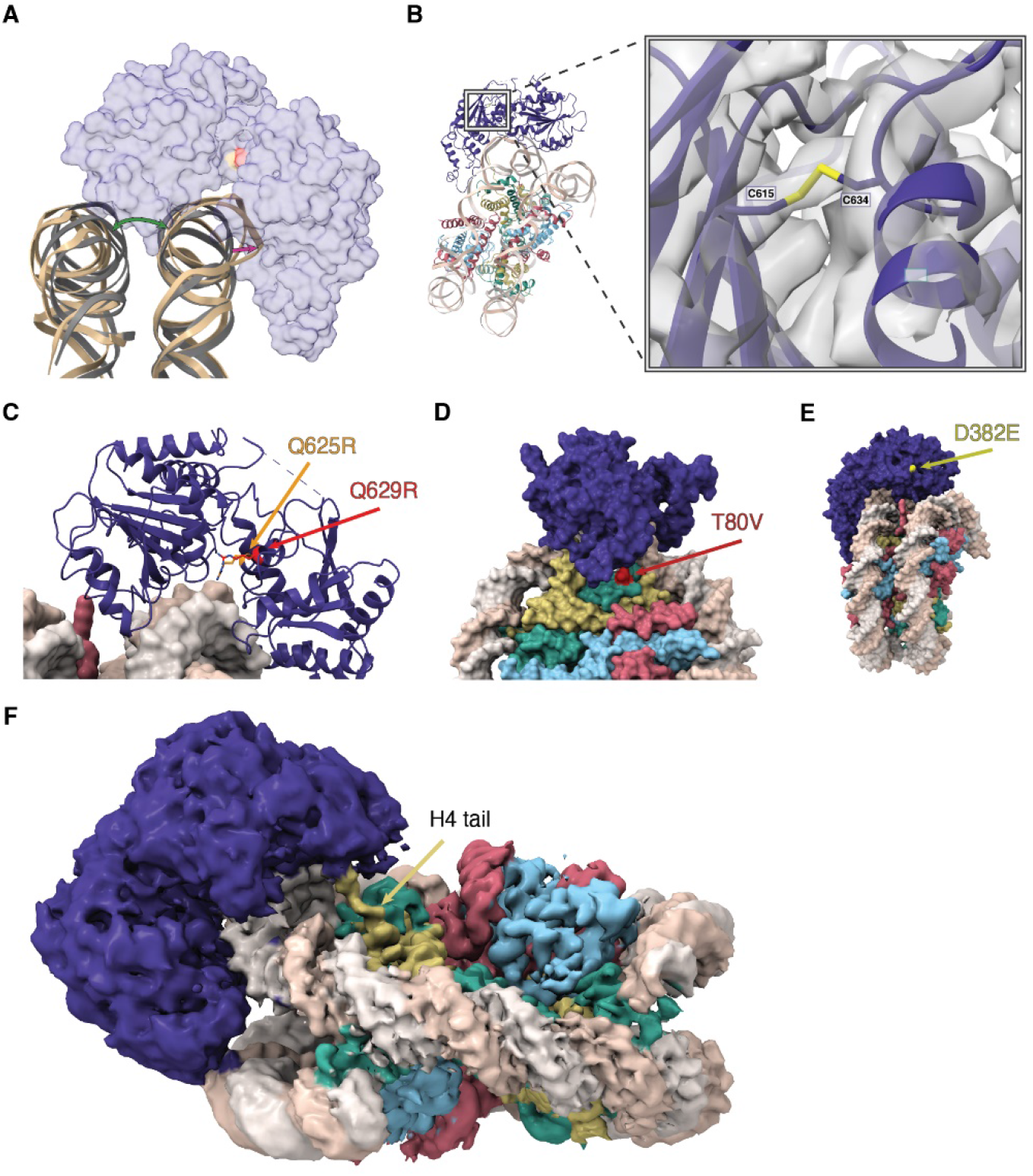
Structure and function of the Helicase C terminal and ATPase lobes. (A) A cartoon view of the DNA distortion caused by DDM1 binding. The DNA backbone of the DDM1- nucleosome model (tan) was aligned to a naked nucleosome DNA backbone (grey, PDB code 1KX5), showing the distortion of DNA where DDM1 is bound to the nucleosome, as well as distortion on the other gyre. A transparent surface model of DDM1 is shown for clarity. The green arrow shows the distortion (opening) of the gyre position caused by DDM1 binding to the nucleosome, and the magenta arrow represents the distortion of the DNA backbone. (B) A view of the disulfide bond formed between C615 and C634, connecting two regions in the HELICc domain of DDM1. The molecular model is shown as ribbons, cryo-EM density is shown as a gray volume. The first mutation of *ddm1* to be isolated, *ddm1-1*, substitutes C615 for Y and has a strong DNA methylation defect. (C) Highly conserved glutamine residues Q625 (red) and Q629 (orange) project between the lobes and are mutated to arginine in human HELLS (identified in ICF syndrome proband E) and in Arabidopsis *ddm1-9* (where it results in hypomethylation), respectively. The arginine residues are predicted to contact phosphates in the DNA minor groove and are also highlighted in (A). (D) A surface representation of the DDM1-nucleosome complex, showing the surface exposed D382E mutation that results in a hypomethylation phenotype in *ddm1-14*. (E) A surface representation of the DDM1-nucleosome complex, highlighting the T80V mutation found in the male germline specific histone H3.3 MGH3 (red). T80 directly contacts DDM1 (Fig. 3C inset). (F) A zoomed in view of the refined cryoEM density map, showing the N-terminal tail of histone H4 extending into the density observed for the DEXD ATPase domain of DDM1. Color coding of histone variants and DDM1 as in Fig.3.

Surprisingly, a disulfide bond was observed in the HELICc domain, bridging C615 and C634 (Figure 4B). Intriguingly, the first allele of *ddm1* to be discovered, *ddm1-1*, has a C615Y substitution predicted to specifically disrupt the S-S bond (Figure S4B), and has strong defects in DNA methylation and DNA repair ^49, 69^. A second allele (*ddm1-9*) has strong silencing defects, and lies between the two cysteines in an absolutely conserved glutamine found in all Snf2 remodelers (Q629R) that lies in the cleft between the two lobes (Figure 4C) ^70^. Although this arginine is too far to contact DNA in the nucleotide-free open conformation (Figure S3A), ADP bound structures of other Snf2 remodelers (Figure S3B) indicate a closed conformation and potential DNA contact ^68^. Intriguingly, a substitution in HELLS, identified in an ICF patient, occurs in a second conserved glutamine nearby (Q625R in DDM1 alignment) indicating conservation of function in humans (Figure 4C) ^27^. In both cases, arginine substitutions are predicted to contact the phosphate backbone around the site of DNA distortion (Figure 4C). A third allele of *ddm1* with strong silencing defects (*ddm1-14*) has a surface substitution (D382E) potentially involved in interacting with protein partners (Figure 4D) ^70^. This surface has a high degree of conservation with other Snf2 remodelers (Figure S3C).

The HELICc domain of DDM1 interacts with histones H3.3 and H4. The majority of the interface is formed by a loop in DDM1 (residues 480-487) that makes contact with H3 at the C-terminus of its α1 helix (residues 73-81) (Figure 3C). In this region, DDM1, DNA, histone H3.3, and histone H4 all contact one another. With histone H3.3, DDM1 makes direct contact at residue T80 (Figure 3C). Intriguingly this residue is substituted by V in the male germline specific variant MGH3 (Figure 4E; Figure S4A), which has been implicated in epigenetic inheritance ^54^ and is mislocalized in heterochromatin in pollen from *ddm1/+* plants (Figure 1C,E). A full experimental density map reveals that additionally, the N-terminal tail of histone H4 extends into a pocket of DDM1, possibly serving as another site for histone mark recognition, although the side chains cannot be resolved (Figure 4F). The H4 tail extends into the same domain in other remodeler structures, including Snf2h and Snf2, but takes a different direction at approximately residue 22 toward the N-terminal end. In the case of DDM1, an aromatic cage comprised of up to 3 tyrosine residues in DDM1 appears at the base of the unstructured H4 tail (Figure S3E). Since the first structured residue is at position 21, the aromatic cage could potentially interact with lysine-20 if it were methylated (Figure S3E). Only one of the three aromatic residues is conserved in HELLS: the positions of key residues in DDM1 and HELLS are summarized in Figure S4B.

### DDM1 has an N-terminal autoinhibitory domain and has remodeling specificity for histone variants

While the ATPase and HELICc domains of DDM1 are well-conserved with Snf2 and well-structured, the N-terminal domain is predicted to be unstructured (Figure 5A; Figure S4C). To establish the role of this N-terminal domain in DDM1 function, we expressed and purified various recombinant DDM1 proteins in *E. coli* and subjected them to peptide binding and ATPase activity assays. Microscale thermophoresis (MST) affinity assays revealed binding of full length DDM1 with unmodified H4 peptides (Figure 5B). Modified H4K20me1, H4K20me2 and H4K20me3 peptides had similar affinities *in vitro*, while fully acetylated H4K5K8K12K16Ac peptides failed to bind (Figure 5B). Recombinant DDM1 has previously been shown to have DNA-dependent ATPase and nucleosome sliding activity, but only when expressed in insect cells and not in *E.coli* ^26^. Consistently, we found only low levels of DNA dependent ATPase activity for full length recombinant DDM1 from *E. coli*, which were further reduced by disruption of the disulfide bond by C615S mutation (Figure 5C). Importantly, however, we found much higher levels of ATPase activity for N-terminally truncated DDM1(Δ1-132) indicating the presence of an N-terminal autoinhibitory domain (Figure 5C). By analogy with ISWI, we named this autoinhibitory domain DDM1 AutoN (Figure 5A). H4 peptide binding assays indicated a slightly higher affinity for the N-terminal truncation that lacks AutoN (Figure 5B).

**Figure 5.**
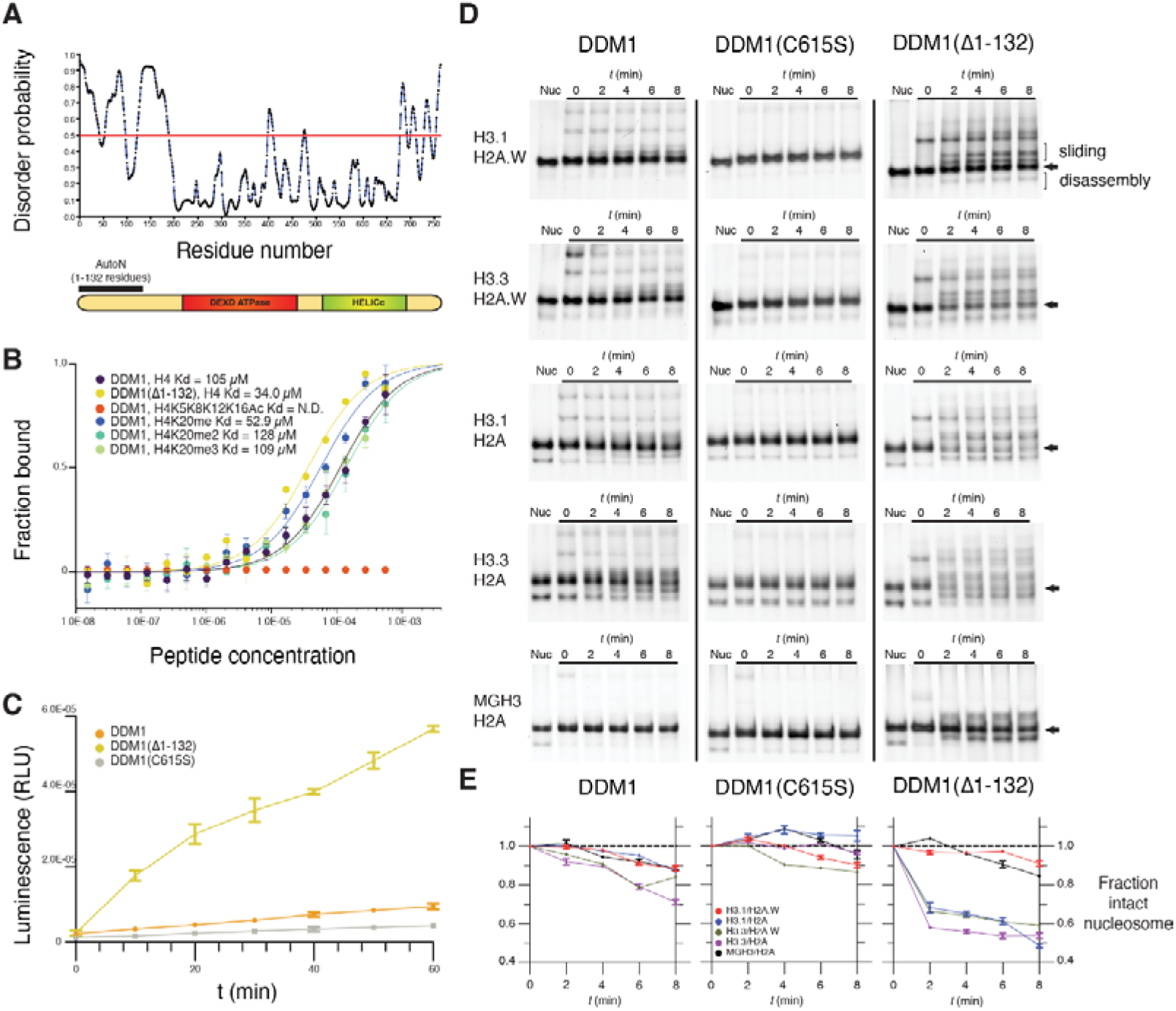
An N-terminal autoinhibitory domain regulates H4 peptide-binding, ATPase and nucleosome remodeling activities of DDM1 with histone variants. (A) Disorder predictions for DDM1 were calculated with PrDOS95. The red line indicates the threshold corresponding to a false positive rate of 5%. The autoinhibitory domain (1-132 residues) AutoN is indicated in the diagram along with DEXD ATPase and HELICc domains. (B) Binding affinities between DDM1 and H4 peptides shown as the fraction bound at peptide concentrations measured by microscale thermophoresis (MST). KD values were estimated by fitting algorithms provided by the supplier (Methods). Binding was not detected (N.D.) for H4K5K8K12K16Ac (quadruple acetylation), but was detected for unacetylated H4, H4K20me1, H4K20me2, and H4K20me3 peptides. Truncation of AutoN in the DDM1(Δ1-132) enzyme resulted in higher binding affinity consistent with AutoN competing with the H4 tail. (C) DNA-dependent ATPase activities for recombinant DDM1, DDM1(Δ1-132) and DDM1 C615S. ATPase activities are given as luminescence with relative light units (RLU). The X-axis indicates ATPase reaction time. Error bars represent standard deviations from two independent replicates. The relative rate enhancement for ATPase activity between DDM1(Δ1-132) and DDM1 is 6.4x. DDM1C615S has a further reduction of 2.3x relative to DDM1. (D) Nucleosome remodeling assays with 0N60 mono-nucleosomes (147bp Widom 601 DNA plus 60bp linker) were performed with octamers of H2B, H4 and combinations of H3 and H2A variants as shown. Center-positioned nucleosomes (arrows) were incubated with DDM1, DDM1 C615S, or DDM1(Δ1-132) at t=0 mins, and then remodeled upon addition of ATP by sliding (slower migration) and disassembly (faster migration). (E) Quantification of remodeling activities for DDM1, DDM1 C615S and DDM1(Δ1-132) are shown below each assay series as the fraction of intact nucleosomes (arrows) remaining at each timepoint relative to t=0. Error bars indicate standard deviation, and are too small to be resolved for H3.3H2A.W. DDM1 C615S had little or no remodeling activity and was used as a control.

Next, we performed nucleosome remodeling assays using nucleosomes composed of H2A, H2A.W, H3.1, and H3.3 variants and the Widom 601 147bp DNA fragment plus 60bp linker (Figures 5D,E). In these assays, nucleosomes come to equilibrium at the center of the fragment, or at the ends, but ATP-dependent DNA translocation activity can ”slide” them resulting in altered mobility on a native gel. ATP-dependent unwrapping activity, on the other hand, can disassemble the octamer into hexasomes and tetrasomes, or remove the octamer altogether ^19^. After incubating nucleosomes with DDM1, DDM1C615S, or DDM1(Δ1-132) at t=0, ATP was added and samples taken every 2 minutes. As an important control, ATP-dependent remodeling was not observed with the DDM1 C615S catalytic mutant enzyme on any of the variant combinations (Figure 5D). The N-terminal truncated form of DDM1, on the other hand, had strong remodeling activity on most histone variant combinations (Figure 5D). In contrast, full-length DDM1 had only weak activity on H3.1-containing nucleosomes, as previously reported ^26^ but stronger activity on H3.3 nucleosomes (Figure 5D).

DDM1 remodeling activity resulted in a pronounced ATP-dependent reduction in intact nucleosomes over time. The proportion of intact nucleosomes (arrows, Figure 5D) were quantified by comparison with levels at t=0 in each reaction. For full length DDM1, reduction of intact nucleosomes was only observed for H3.3 containing nucleosomes (green and purple lines, Figure 5E). For the truncated enzyme, remodeling was much more pronounced, but reduction of intact nucleosomes was restricted to nucleosomes containing H3.1/H2A, H3.3/H2A or H3.3/H2A.W, reaching equilibrium at roughly 50% intact nucleosomes (Figure 5E). Despite close similarity of H3.1 and H3.3 (Figure S4A), DDM1 could not reduce the level of intact nucleosomes composed of both H3.1 and H2A.W (red lines, Figure 5E), despite the presence of slower migrating bands indicating potential sliding activity. The germline specific H3.3 variant MGH3 differs from H3.3 at 12 positions, 4 of which are in the globular domain, one of which (Y41F) is shared with H3.1 (Figure S4A) and one of which (T80V) contacts DDM1 directly (Figure 3C; Figure 4E). We performed remodeling assays on MGH3 variant nucleosomes, and found that replacing H3.3 by MGH3 made H2A nucleosomes resistant to remodeling by DDM1, differing dramatically from both H3.1 H2A and H3.3 H2A nucleosomes (Figure 5D), and resembling H3.1 H2A.W nucleosomes instead (Figure 5E).

To further test the idea that remodeling of histone H3 variants underlies the role of DDM1 in epigenetic inheritance, we performed confocal microscopy and live imaging of cycling root tip cells, to examine the localization of DDM1 and histone variants. We found that DDM1-mCherry colocalized with H3.1-CFP at chromocenters during S phase (Figure 6A) ^39, 71^ when deposition of H3.1 is mediated by CAF-1 ^72^. In most interphase cells, however, DDM1-GFP was diffusely localized in the nucleoplasm along with H3.3-RFP (Figure 6B). In live imaging experiments, DDM1 was recruited to chromatin in G1, remained until G2, and then dissociated upon mitosis (Videos S1 and S2). In humans, HELLS also has diffuse nucleoplasmic localization, but mutation of the Walker-A ATP binding site (K237Q) results in tight association with chromocenters and reduced soluble fractionation in nuclear extracts ^73^. Given the high conservation between LSH/HELLS and DDM1 in this region (Figure 6C), we generated an equivalent DDM1^K233Q^ mutation in the pDDM1:DDM1-mCherry transgene. Relative to the wild-type transgene fusion, this mutated version of DDM1 also displayed enhanced chromocenter localization in WT cells (Figure 6C), and reduced partitioning into the soluble nuclear fraction (Figure 6D).

**Figure 6.**
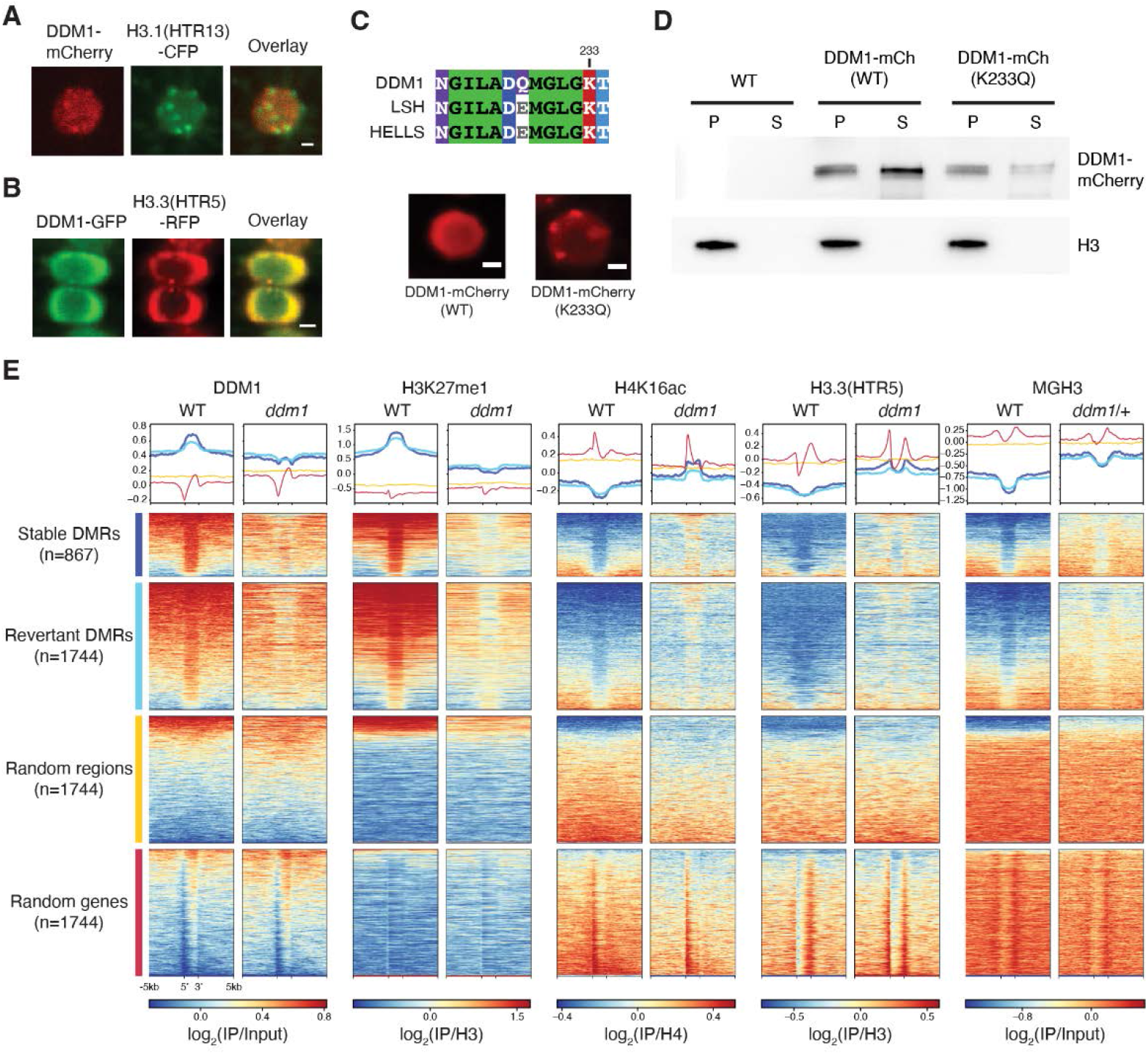
DDM1 remodels H3.3 and H3.1 in interphase at differentially methylated targets of DDM1. (A) Subnuclear localization of DDM1-mCherry in root tip cells as compared to H3.1(HTR13)-CFP during presumptive late S-phase (marked by H3.1-labeled chromocenters). (B) Subnuclear localization of DDM1-GFP as compared to H3.3(HTR5-RFP) during interphase. The scale bar indicates 2 µm. See also Supplemental Videos S1and S2. (C) Conserved amino acids of the ATP binding sites for DDM1 and orthologs, LSH (mouse) and HELLS (human). K233Q Walker A mutation disrupts ATP binding and causes enhanced chromocenter localization of DDM1-mCherry fusion in a WT background. Scale bar indicates 2 µm. (D) Western blot of DDM1- mCherry (DDM1-mCh) for the wild-type (WT) and mutant form (K233Q) from soluble (S) and chromatin/pellet (P) fractions indicates failure to release catalytic mutant from chromatin. H3 was used as loading control. Non-transgenic WT was used as negative control. (E) Comparisons of DDM1, H3K27me1, H4K16ac and H3.3(HTR5) chromatin association by ChIP-seq in WT and *ddm1* leaves, as well as MGH3 in pollen from WT and *ddm1*/+ plants. Stable and revertant differentially methylated regions (DMRs) lost DNA methylation in *ddm1* mutants, and stable DMRs never regained methylation when DDM1 was reintroduced ^74^. Thus, DMRs represent epigenetic targets of DDM1. Heatmaps and metaplots were generated using DeepTools ^95^, where each region was scaled to 2kb with 5kb upstream and 5kb downstream with a binsize of 10bp, and sorted based on DDM1 levels in WT. Metaplots above each heatmap show the mean value for each region. Random regions are revertant DMRs reshuffled randomly in the genome, whereas random genes correspond to the same number of protein coding genes selected at random (see Table S2). DDM1 and H3.1 (H3K27me1) are specifically enriched at DDM1 targets (DMRs), while H4K16Ac, H3.3 and MGH3 are specifically depleted, except in *ddm1* mutants. Similar analysis was performed on all transposable elements for comparison (Figure S5).

### The role of DDM1 in epigenetic inheritance during the cell cycle

In serial crosses to wild-type, unmethylated DNA is epigenetically inherited from *ddm1* mutants, and differentially methylated regions (DMRs) between WT and *ddm1* can be divided into two classes depending on their inheritance ^74^. The first class are stable DMRs that are never remethylated in serial crosses to WT plants. The second class of DMRs eventually revert to WT in similar crosses, although reversion can take multiple generations. Both classes of DMRs represent epigenetically inherited targets of DDM1 and are almost exclusively comprised of transposable elements ^74^. We mapped our ChIP-seq reads to these DMRs, and compared them to random sequences and genes (Figure 6E). Additionally, we performed ChIP-seq using antibodies for H4K16ac, a highly conserved modification that marks active chromatin and has been shown to accumulate in chromocenters in *ddm1* mutants ^75^. H4K16ac is a reliable marker for histone H4 acetylation in plants, and is accompanied by H4K12, K8 and K5 acetylation ^76^, which we found prevents binding of DDM1 to H4 tails *in vitro* (Figure 5B). Using these DMRs as proxies for DDM1 activity, we first found that DDM1 is strongly enriched precisely over these DMRs, consistent with DMRs being DDM1 targets (Figure 6E), and mostly comprising pericentromeric transposable elements (Figure S5A,B). Similarly, H3K27me1 was also strongly enriched in DDM1 targets but depleted in *ddm1*. In contrast H3.3, MGH3 and H4K16Ac are precisely excluded from DDM1 targets in WT plants, but encroach into these regions in *ddm1* mutants (Figure 6E; Figure S5). Notably, H4K16Ac was especially enriched in stably inherited DMRs in *ddm1* mutants. Thus, histone H3.3, MGH3 and H4K16Ac are each strongly anti-correlated with both DDM1 and H3.1, specifically at the epigenetically inherited and differentially methylated targets of DDM1.

## Discussion

Nucleosome remodeling is an important pre-requisite for DNA metabolism, including DNA replication, repair, recombination, and transcription ^19^. Here we show that DDM1 promotes DNA methylation by preferentially remodeling heterochromatic nucleosomes during S phase, when H3.1 and H2A are deposited, allowing DNA methylation of CG dinucleotides by the methyltransferase MET1 (Figure 7). Subsequent incorporation of H2A.W stabilizes H3.1, but not H3.3 nucleosomes, allowing DNA methylation of non-CG cytosines in G2 by the chromomethylases CMT3 and CMT2, which depends on lysine 9 di-methylation of intact histone H3 nucleosomes ^77^. DDM1 is then evicted in mitosis.

**Figure 7.**
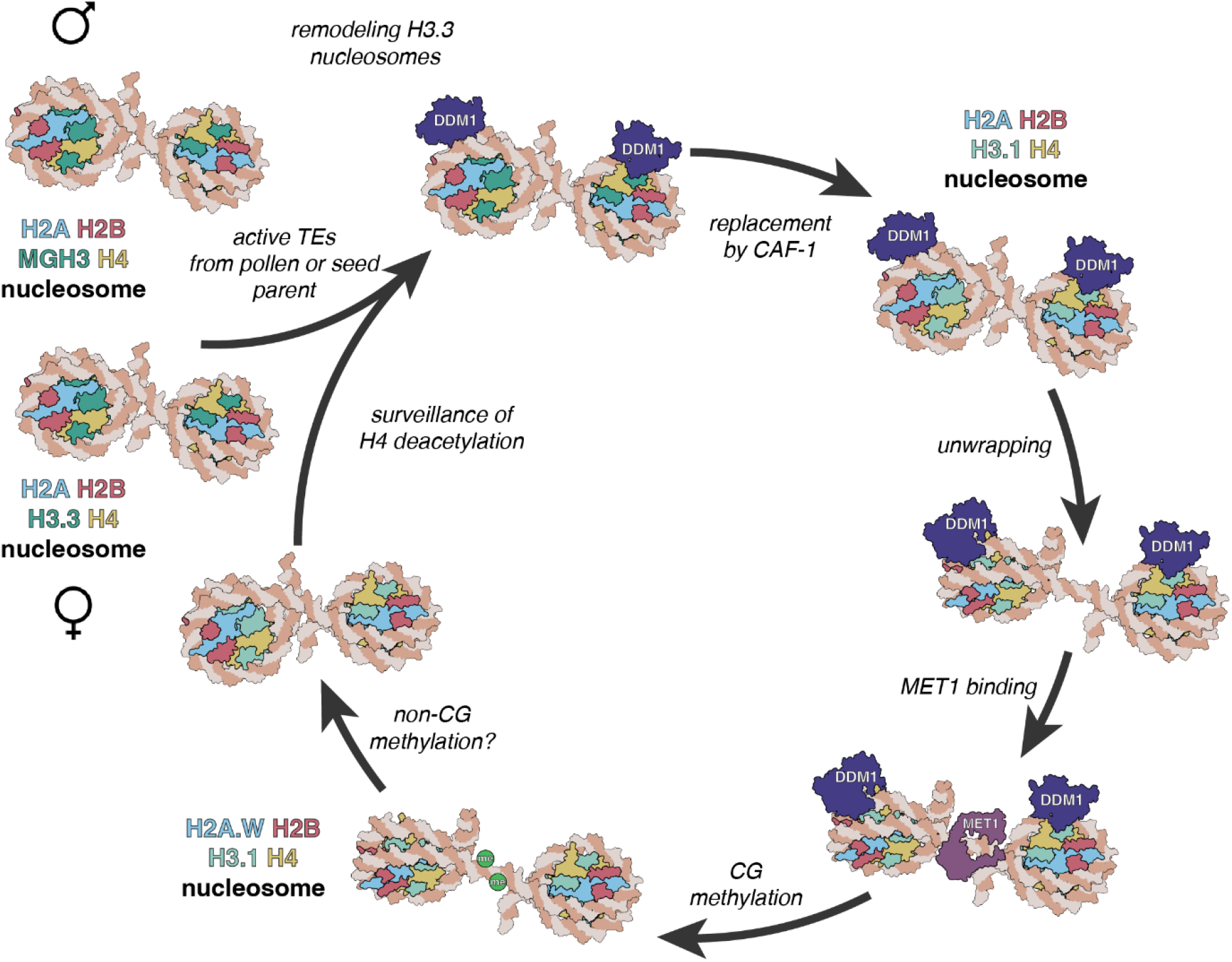
A model for epigenetic inheritance of unmethylated transposons from *ddm1* mutants. Active transposable elements (TEs) from male and female gametes are unmethylated and comprise MGH3 H2A and H3.3 H2A nucleosomes, respectively. In the zygote, H3.3 H2A nucleosomes are remodeled by DDM1 before replication, allowing deposition of H3.1 in S phase by CAF-1. Unwrapping of H3.1 H2A by DDM1 permits access to the methyltransferase MET1 allowing CG methylation. Subsequent incorporation of H2A.W stabilizes the nucleosome, possibly promoting H3K9 di-methylation and CHG methylation by the chromomethylase CMT3 (not shown). MGH3 H2A nucleosomes, inherited from pollen, are resistant to remodeling after fertilization. They are eventually replaced in the embryo by H3.3 H2A nucleosomes, but acetylation of histone H4 and other marks of active euchromatin prevent recognition by DDM1.

Mutants in H3.1 and H3.3, their chaperones, and their modifying enzymes, all exhibit strong genetic interactions with mutants in *ddm1,* supporting the essential role of this mechanism in heterochromatic DNA replication ^42^ and DNA methylation ^2–4^.

Remodeling assays with variant histones supported our conclusions. Octamers containing both H3.1 and H2A.W were much more stable in remodeling assays with DDM1 than other octamers (Figure 5D,E), although slower migrating nucleosomes were evidence of sliding or unwrapping activity, as opposed to disassembly. These differential activities could result in the preferential deposition of H3.1 and H2A.W by destabilizing nucleosomes containing H3.3 or H2A ^34, 78^. This mechanism may also account for the dependence on DDM1 for deposition of histone H1 ^34, 35^ and for the requirement of LSH for histone macroH2A deposition *in vivo* ^37^ and macroH2A exchange *in vitro* ^36^. This is because assembly of H1 linkers, and of H2A/H2B dimers, depend on previous assembly of H3/H4 tetramers ^19^. In the presence of DDM1, histone gene mutations indicate that histone H1 inhibits DNA methylation ^29, 32–35, 37^, while histone H3.3 and histone H2A.W can actually promote it ^55^, although the effects are small. In the absence of remodeling by DDM1, access to DNA methyltransferase is blocked by the residual H3.3 nucleosomes. What then is the role of H2A.W or macroH2A? We identified an N-terminal autoinhibitory domain in DDM1 that is analogous to the N-terminal intrinsic autoinhibitory domain (AutoN) of ISWI ^79^. ISWI AutoN inhibits ATP hydrolysis but this inhibition is relieved upon interaction with an acidic patch on H2A ^80^. In a recent study, the N-terminal domain of DDM1 was shown to bind H2A.W in a similar way ^34^, which could promote ATPase activity. Thus H3.1H2A.W sliding activities we observe might promote DNA methylation of intact nucleosomes in the G2 phase of the cell cycle by CMT2 and CMT3 ^77^, accounting for the modest reductions in non-CG methylation observed in *h2a.w* mutants ^34^.

Single-particle cryo-EM reconstruction of DDM1 bound to variant nucleosomes supports this model, in that DDM1 engages the nucleosome at SHL-2 making direct contact with histone H3 at T80 and D81, just 6 amino acids from H87, the residue which confers specificity to histone H3.3 nucleosome assembly in *Arabidopsis* (Figure S4A) ^81^. The H3- interacting loop of DDM1 is only moderately conserved between plants and mammals (Figure S4B), which have differing variant residues in histone H3.3, indicating co-evolution with histone H3.3. A disulfide bridge in the C terminal helicase domain is required for DNA-dependent ATPase activity and chromatin remodeling *in vitro*, and is disrupted in the very first *ddm1* allele (*ddm1-1*), providing strong support that ATP-dependent remodeling of histone variants is required for DNA methylation (Figure 4B; Figure 5C,D,E). In mammalian cells, histone residues H3K79 and T80 are modified by methylation and phosphorylation, respectively, during mitosis, which might be expected to prevent interaction with DDM1 ^82^. Consistently, imaging of wild-type and catalytic mutant DDM1 revealed colocalization with H3.3 in interphase, and with H3.1 during S phase, but DDM1 was lost during mitosis when histone H3.3 accumulates instead (Figure 6; Videos S1 and S2). The reverse is true for ATRX, which deposits H3.3 in heterochromatin in mammals ^58–61^, and removes macroH2A ^83^ likely promoting heterochromatic transcription in G1 ^84^. In *Arabidopsis*, ATRX is also required for H3.3 deposition in heterochromatin ^62, 85^ and may have a reciprocal function to DDM1 in mitosis (Graphical abstract). Intriguingly, we found that ATRX is also required for DNA methylation of a subset of DDM1 targets, confirming that remodeling itself, rather than specific histone variants, is required for DNA methylation ^29^.

The cryo-EM single particle reconstruction has also revealed important clues to heterochromatic specificity. The nucleosomal contacts made with histone H4 by DDM1 are similar to those made by ISWI/Snf2h ^63, 66, 86^. Histone H4 deacetylation is a hallmark of heterochromatin, and in ISWI the H4 tail must be deacetylated to stimulate remodeling activity ^19, 79^. Chd1 also interacts with the H4 tail for DNA translocation activity ^67^ and physically associates with HIRA for H3.3 deposition ^23^. Binding assays with DDM1 demonstrated specific affinity for deacetylated H4 tails but not for fully acetylated H4 peptides (Figure 5B), consistent with specific activity for heterochromatic nucleosomes which are strongly deacetylated in Arabidopsis (Figure S5B). ISWI AutoN competes with the H4 tail, and DDM1 without AutoN had higher affinity for H4 peptides (Figure 5B), consistent with this mechanism. In striking agreement with these conclusions, H4K16 acetylation, which is a reliable marker for H4 acetylation, is enriched in precisely those regions of heterochromatin in which DDM1 is depleted (Figure 6D, Figure S6A,B). DDM1 also has high affinity for H4K20me1, H4K20me2, and H4K20me3 peptides, and although their association with plant heterochromatin is less well established, they might provide specificity for other roles of DDM1, such as DNA repair.

Our results shed important light on the role of DDM1 in epigenetic inheritance. We found that DDM1 interacts with MET1 in Arabidopsis (Figures S1D,E), similar to the interaction of LSH with DNMT1 in the mouse ^10, 50, 51^. DDM1 remodeling activity allows chromatin access to MET1 ^29^, but *MET1* is also required for DDM1 binding to chromatin (Figure S1F). Histone H4 deacetylation is likely responsible, as it depends on *MET1* ^87, 88^. Unmethylated DNA is epigenetically inherited for multiple generations from *ddm1*, *met1* and *hda6* mutants, but not from other histone and DNA modification mutants ^89, 90^, consistent with this circular logic. However, it has been previously reported that H4K16ac is restored to wild type levels when DDM1 activity is restored in heterozygotes ^75^, as is H2A.W ^34^, arguing against a direct role in epigenetic inheritance. Instead, we found that the Male Germline H3.3 variant MGH3 is mislocalized to heterochromatin at the nuclear periphery in sperm cells from *ddm1/+* and *met1/+,* even when wild-type DDM1 function was restored (Figure 1E; Figure 6E). MGH3 has a T80V substitution predicted to alter interaction with DDM1 (Figure 4E; Figure S4A), potentially accounting for failure to remove ectopic MGH3 in WT pollen. Consistent with this idea, remodeling assays with DDM1 revealed that MGH3 H2A nucleosomes were much more resistant to remodeling than H3.3 H2A or H3.1 H2A nucleosomes. While the precise mechanism remains to be determined, interaction with histone H3 variants *in vivo* and *in vitro* suggests that they contribute to silencing by DDM1.

These results lead to a model for epigenetic inheritance of unmethylated transposable elements that depends on differential nucleosome remodeling (Figure 7). Active transposons are transcribed, and associated with H3.3 and H2A. When active transposons are introduced into WT plants by genetic crosses with *ddm1* mutants, DDM1 remodels chromatin by unwrapping H3.3 and H2A nucleosomes, promoting the assembly of H3.1 H2A.W nucleosomes instead. Remodeling promotes access to DNA methyltransferases, DNA methylation and silencing. An exception occurs if active transposons are introduced from the male, because MGH3 replaces H3.3 in sperm cells, where it cannot be removed by DDM1. Subsequent replacement in the zygote with H3.3, and H4K16 acetylation results in epigenetic inheritance of the active transposon. One prediction of this model is that transgenerational epigenetic inheritance of unmethylated transposons should occur preferentially through the male germline. Remarkably inheritance of active transposons is indeed paternally biased ^91^ resulting in the gradual restoration of DNA methylation from one seed generation to the next ^48^. We previously found that MGH3 is also resistant to H3K27 tri-methylation by Polycomb repressor complex ^54^, resembling placeholder nucleosomes that underlie paternal inheritance of DNA methylation in zebrafish ^92^. Thus, chromatin remodeling of histone variants by DDM1 underlies the epigenetic inheritance of DNA methylation in plants. Transgenerational inheritance is much less common in mammals ^93^, but close parallels with HELLS and Lsh suggest similar mechanisms may operate in the mammalian germline as well. A recent study has indicated that transgenerational inheritance of a methylated promoter in the mouse occurs despite the reprogramming and loss of DNA methylation in the embryo, suggesting a similar placeholder mechanism may be at work in mammals as well ^94^.

## Acknowledgments

We thank Crisanto Gutierrez for providing H3.1-GFP and H3.3-RFP *Arabidopsis* reporter lines and Frédéric Berger for providing the *htr4 htr5 htr8/+* and HTR13-CFP lines. We thank Eric Richards, Vincent Colot and Tetsuji Kakutani for discussions on the genetics of *ddm1*, and Cigall Kadoch, Filipe Borges and Jean-Sebastien Parent for helpful advice. We thank Joe Calarco (*in memoriam*) for the anti-DDM1 polyclonal antibodies. We acknowledge technical assistance for live imaging from Nour El-Amine, and informatics help from Leandro Quadrana (Ecole Normale Superieur, France). Research in the Martienssen laboratory is supported by the U.S. National Institutes of Health (NIH) grant R01 GM067014, the National Science Foundation Plant Genome Research Program, and the Howard Hughes Medical Institute. The authors acknowledge assistance from the Cold Spring Harbor Laboratory Shared Resources, which are funded in part by a Cancer Center Support grant (5PP30CA045508). This work was also supported by Grant #R35GM128661 from the National Institutes of Health to Y.J., the Wellcome Trust [104175/Z/14/Z], Sir Henry Dale Fellowship to P.V. and through funding from the European Research Council (ERC) under the European Union’s Horizon 2020 research and innovation program (ERC-STG grant agreement No. 639253 to P.V.). The Wellcome Centre for Cell Biology is supported by core funding from the Wellcome Trust [203149]. We are grateful to the Edinburgh Protein Production Facility (EPPF) for their support. The EPPF was supported by the Wellcome Trust through a Multi-User Equipment grant [101527/Z/13/Z]. LJ and RAM are Investigators of the Howard Hughes Medical Institute.

## Author contributions

SCL, LJ and RAM designed the study; SCL, JJI, DWA, JL, HK, UR, CL, UR, SB, and VM performed the experiments; JC, SCL, JJI, DWA, BB, DG, YJ, LJ, and RAM analyzed the data and/or its significance; SCL and RAM wrote the manuscript with contributions from JC, JJI and PV. YJ, PV, LJ, and RAM acquired funding.

## Competing interests

The authors declare no competing interests.

## STAR Methods

### Resource availability

#### Lead Contact

Further information and requests for resources and materials should be directed to and will be fulfilled by the lead contact, Robert A. Martienssen.

#### Materials availability

All materials generated in this study are available upon request to the lead contact.

#### Data and code availability

- ChIP-sequencing and Bisulfite-sequencing data have been deposited at GEO and are publicly available as of the date of publication. Accession numbers are listed in the key resources table (GEO study: GSE231563).
- This paper also analyzes existing, publicly available data. These accession numbers for the datasets are listed in the key resources table.
- Coordinates and the cryo-electron microscopy map for the DDM1-nucleosome complex have been deposited in the Protein Data Bank and are publicly available as of the date of publication. Accession numbers are listed in the key resources table. (PDB: 7UX9; EMD-26855).
- All original code has been deposited at https://github.com/martienssenlab/DDM1-manuscript
- Any additional information required to reanalyze the data reported in this paper is available from the lead contact upon request.

### Experimental model

#### Seed stocks and plant materials

Plants were grown under long day conditions at 22°C. Seeds were grown on ½ Murashige and Skoog (MS) medium and seedlings were transplanted to soil 7 day after germination for BS-seq, or harvested at 10 days for ChIP. For MGH3, ChIP was performed on mature pollen grains.

All genotypes including wild-type and *met1-1*, *met1-7*, *cmt3-11*, *ddm1-2*, *ddm1-10*, *fas2- 4*, *hira-1*, and *atrx-2* mutants are in the Col-0 background and described in Table S3. The pHTR3:HTR3-GFP (referred herein as H3.1-GFP) and pHTR5:HTR5-RFP (referred to as H3.3-RFP) lines were previously reported ^39^, as were *htr4 htr5 htr8*/+ ^55^, pMGH3:MGH3- GFP ^103^, pHTR13:HTR13-CFP ^104^ and pDDM1:DDM1-GFP ^44^. To generate mCherry reporter lines, genomic fragments of DDM1 (AT5G66750) and MET1 (AT5G49160) were cloned into the vector pDONR221 with the mCherry fragment inserted before the stop codon by NEBuilder HiFi DNA Assembly Master Mix (New England Biolabs) (Table S4 for primer sequences). mCherry constructs in pDONR221 were transferred into the pB7WG binary destination vector using Gateway LR Clonase II (Thermo Fisher Scientific). pDDM1:DDM1^K233Q^-mCherry was generated from pB7WG-pDDM1:DDM1-mCherry using the Q5 Site-Directed Mutagenesis Kit (New England Biolabs). For biomolecular complementation experiments, DDM1 and MET1 coding sequences were cloned into pBiFC2 and pBiFC4, respectively. DDM1-GFP constructs were shown to complement *ddm1* mutants previously ^44^. *met1-1* mutants were late flowering (57 leaves, n=23), unlike WT Col-0 (15.6 leaves, n=12) and *met1-1* mutants were partially complemented by MET1-mCherry (37.5 leaves, n=4). *ddm1 fas2/+* plants had 19% aborted seed (n=122) while *fas2 ddm1/+* plants had 17% aborted seeds (n=75). No double mutants were recovered (n>150).

### Methods

#### Chromatin immunoprecipitation (ChIP)

ChIP was performed as previously described ^105^, starting with 1g of seedlings. In brief, after crosslinking in 1% formaldehyde for 10 min, the tissue was ground to a fine powder in liquid nitrogen, chromatin was extracted with 1% SDS Tris-based lysis buffer and sonicated to ∼200bp fragments. Chromatin was cleared with protein A magnetic beads and incubated overnight with the antibody listed below. Immune complexes were eluted with low, and then high salt buffers, before reversing crosslinks and purifying DNA fragments with ChIP DNA clean and concentrator kit (Zymo Research). Low input ChIP for MGH3-GFP was performed with ∼10e8 pollen grains. Fixation was for 15 min and subsequent grinding with acid-washed glass beads. DNA fragments underwent a preliminary phenol-chloroform and ethanol precipitation step before clean up.

For histone modifications, anti-H3K27me1 (Active Motif; 61015) and anti-H3 antibodies (Abcam; ab1791), or anti-H4K16ac (EMD millipore; 07-329) and anti-H4 (abcam; ab10158) antibodies were used. For ChIP experiments against H3.1-GFP, H3.3-RFP, and DDM1-mCherry, GFP-trap (ChromoTek; gtma-10) or RFP-trap (ChromoTek; rtma- 10) magnetic beads were used. For MGH3-GFP ChIP, anti-GFP (Thermo Fisher scientific; A-11122) polyclonal antibody was used. Polyclonal DDM1 antibodies were raised against a synthetic peptide (TINGIESESQKAEPEKTGRGRKRKAASQYNNTKAKRAVAAMISRSKE) outside the SWI/SNF domain by Covance antibody services. Anti-H3 was used as control for H3K27me1 and H3.3, anti-H4 was used as control for H4K16ac, and input DNA were used as controls for MGH3 and DDM1 ChIP-seq. After ChIP, qPCR was performed using KAPA SYBR FAST qPCR Master Mix (Kapa Biosystems). qPCR primers are listed in Table S4. ChIP-sequencing (ChIP-seq) libraries were prepared with ∼200 bp insert size using NEBNext Ultra II DNA Library Prep Kit for Illumina (New England Biolabs). The ChIP-seq libraries were sequenced using an Illumina NextSeq platform with 75-cycle single reads for H3K27me1 and HTR5 samples and with paired-end 151 cycles for DDM1 and H4K16ac samples. MGH3 libraries were prepared by Fasteris SA (Switzerland) and sequenced on HiSeq platform to 100bp paired-end reads for WT and 100bp single reads for ddm1/+. Only read1 was processed from the WT sample to compare to single-end ddm1/+. Sequencing metrics on all ChIP-seq libraries are listed in Table S5. ChIP-seq data were analyzed as previously described ^106^, trimming adapters with cutadapt ^107^ but using bowtie2 ^108^ to map to TAIR10, before filtering primary alignements with samtools^109^. Two independent replicates of ChIP-seq were obtained for each antibody and genotype except for MGH3-GFP, which was compared instead to H3.3. The Pearson correlation coefficient was at least 0.8 for the replicates (Figure S6A). When available, the IP and control of the two biological replicates were merged after mapping and deduplication with samtools, and then the signal tracks (bigwig) were generated with Deeptools ^95^ by calculating the log2 fold change of IP over control after count-per-million normalization. MGH3 and H3.3 ChIP-seq datasets produced in this study have been compared to previously published datasets GSE120664 ^47^ (Figure S6B) and from GSE34840 ^40^ (Figure S6C), respectively. Figures were generated in R (https://www.r-project.org/) using ggplot2 ^110^ and GViz ^111^ packages. Heatmaps were generated with Deeptools.

#### Bisulfite sequencing (BS-seq)

Genomic DNA (gDNA) was extracted from rosette leaves of 3-4 plants for each genotype with Nucleon PhytoPure Genomic DNA Extraction Kits (Cytiva). 1 μg gDNA samples were sheared to an average size of 400 bp using a Covaris S220 focused-ultrasonicator. BS-seq libraries were made using the NEXTFLEX Bisulfite Library Prep Kit (PerkinElmer). The library samples were sequenced as paired-end 101bp reads with an Illumina NextSeq system. Sequencing metrics on all Bisulfite sequencing libraries are listed in Table S6. Adaptors were trimmed using Cutadapt ^107^ and aligned to the TAIR10 reference genome with Bismark ^112^. Duplicate reads were collapsed, and methylation levels at each cytosine were calculated as a ratio of #C / (#C + #T). DMRs were defined using DMRcaller^113^. A minimum difference of 30%, 20%, and 10% was used for DMRs in CG, CHG, and CHH contexts, respectively. Two independent biological replicates were performed for each genotype, showing high reproducibility (Pearson correlation > 0.9, Figure S6D,E). Figures were generated in R (https://www.r-project.org/) using ggplot2 ^110^ and GViz ^111^ packages.

#### Nuclear fractionation

Soluble and insoluble chromatin fractions were obtained as previously described ^114^. Briefly, 0.3 mL of seedling tissues were incubated in 0.6 mL N1 buffer (15 mM Tris-HCl, pH 7.5, 60 mM KCl, 15 mM NaCl, 5 mM MgCl2, 1 mM CaCl2, 1 mM DTT, and 250 mM sucrose, with Complete Mini EDTA-free protease inhibitor and PhosSTOP (Roche)) on ice for 30 mins. After filtering the extract with 30 μm CellTrics filters (Sysmex), nuclei were isolated by centrifugation at 4 °C and 1000g for 10 min. Nuclei were washed twice with N2 buffer (15 mM Tris-HCl, pH 7.5, 60 mM KCl, 15 mM NaCl, 5 mM MgCl2, 1 mM CaCl2, and 1 mM DTT, with Complete Mini EDTA-free protease inhibitor and PhosSTOP) and subsequently incubated with 600 μL of N3 buffer (15 mM Tris-HCl, pH 7.5, 150 mM NaCl, 5 mM MgCl2, 1 mM CaCl2, and 1 mM DTT, with Complete Mini EDTA-free protease inhibitor and PhosSTOP) for 30 min. The samples were centrifuged at 12,800g and 4 °C for 15 min to yield the soluble and insoluble pellet fractions. Soluble fractions were concentrated by vortexing with StrataClean resin (Agilent) for 1 min. Samples were boiled at 95 °C in SDS loading buffer and used for Western blot experiments. Anti-DDM1, anti-RFP (Rockland; 600-401-379) and anti-H3 (Abcam; ab1791) antibodies were used for detection of endogenous DDM1, DDM1-mCherry, and H3, respectively.

#### Microscopy

Immunofluorescence experiments for leaf nuclei were performed using 3-week-old leaves as previously described ^115^. Monoclonal antibodies of anti-H3K27me1 (Active Motif; 61015) were used as primary antibodies and goat anti-mouse Alexa Fluor 488 (Thermo Fisher Scientific; A-10680) was used as secondary antibody. DAPI (2 μg/mL final concentration) was used for nuclei staining and the samples were mounted with Prolong Diamond (Thermo Fisher Scientific). DDM1-GFP, DDM1-mCherry, HTR3-GFP, and HTR5-RFP were observed with a Zeiss 710 confocal microscope. Live imaging data of DDM1-GFP was acquired using a Perkin-Elmer UltraVIEW VoX confocal microscope.

#### Expression and purification of DDM1 protein

His-TEV-DDM1 and His-TEV-DDM1(Δ1-132) were transformed into the *E.coli* strain BL21-CodonPlus (DE3)-RIPL (Agilent) for large-scale expression using standard methods. Briefly, cultures were grown in Terrific Broth media supplemented with appropriate antibiotic(s) at 37°C to a culture density of approximately ODλ=600 nm of 1.2. Cultures were then cooled in an ice water bath for 15 minutes followed by induction of protein expression with 0.5 mM IPTG. Induction proceeded overnight at 16 °C with shaking at 220 rpm. Cells were harvested by centrifugation at 4000g for 30 minutes at 4 °C. The supernatant was discarded and the pelleted cells were taken for protein purification.

For Ni-NTA purification, cell pellets were resuspended in 20 mL lysis buffer (20 mM Tris, pH 8.0, 300 mM NaCl, 5 mM MgCl2, 10% glycerol, 1 mM TCEP, 20 mM imidazole) per liter culture. Protease inhibitors and 0.1% Triton-X were next added to the resuspension and the cells lysed by sonication. Turbo nuclease (Accelagen) was added to the cell lysate (2.5 units per mL of cell resuspension) and the lysate was then clarified by ultracentrifugation at roughly 100,000g for 30 minutes. The soluble supernatant was taken for affinity purification via batch binding with Ni-NTA resin (2 mL of beads per liter culture), pre-equilibrated with lysis buffer. Batch binding was performed for 2-3 hours at 4 °C with gentle agitation. The Ni-NTA beads were then collected by centrifugation at 1000g for 5 minutes, resuspended in lysis buffer, then transferred to a column for further washing and elution. Beads were washed with 20 column volumes of lysis buffer followed by elution of the target protein in lysis buffer supplemented with 100-250 mM imidazole.

To remove the affinity tag, TEV protease was added in a 1:20 mass ratio (protease:DDM1) and incubated overnight at 4 °C. In addition, DTT was added to a final concentration of 10 mM to limit aggregation. Protein was further purified using a HiTrap Heparin HP column (Cytiva/GE Healthcare Life Sciences). Digested protein was first diluted two-fold with low salt buffer (20 mM Tris, pH 8.0, 1 mM DTT) prior to loading the heparin column.

The target protein was then eluted using a 25-75% gradient of high salt buffer (20 mM Tris, pH 8.0, 1 mM DTT, 1 M NaCl) over approximately 50 mL. Peak fractions were assessed by SDS-PAGE then selected and pooled for further purification.

Pooled fractions were concentrated to 500 μL and applied to a Superdex 200 increase 10/300 column (Cytiva/GE Healthcare Life Sciences). The protein was chromatographed over ∼30 mL at a flow rate of 0.6 mL/min in a running buffer of 20 mM Tris, pH 8.0, 300 mM NaCl, 5 mM MgCl2, 1 mM TCEP. Peak fractions were assessed by SDS-PAGE. Fractions with highly-purified protein were concentrated, then taken for enzymatic assays and/or storage. For long-term storage the protein was flash frozen in liquid nitrogen, then kept at -80 °C. Typical yields were 1-2 mg of highly purified protein (>98% pure as assessed by SDS-PAGE) per liter culture.

#### DDM1 ATPase, remodeling, gel shift, and peptide-binding assays

DDM1 ATPase assays were performed in reaction buffer (10 mM Tris pH 7.5, 50 mM NaCl, 10 mM MgCl2, 20% Glycerol) containing various ATP concentrations and quantified using the ADP-Glo MAX Assay (Promega; Catalog No. V7001) as described previously^116^. The double-stranded DNA substrate was prepared by PCR amplification of the Widom 601 DNA sequence described in the remodeling assays below (see Table S4 for primer sequence information). Methylated DNA was amplified by PCR in the presence of 5m- dCTP rather than dCTP (New England Biolabs). Additional experimental procedures followed the manufacturer’s guidelines. Luminescence was quantified using GloMax-Multi+ Detection System (Promega).

DDM1 nucleosome remodeling assays were performed with mononucleosomes. Histone octamers consisting of *Arabidopsis* H2A.W (AT5G59870), H3.3 (At5g10980), H3.1 (At1g09200), H2A (AT3G20670) and H2B (AT3G45980), and *Xenopus* H4 were assembled as described previously ^117^. Briefly, core histones were expressed in *E. coli*, solubilized from inclusion bodies, and purified by sequential anion and cation exchange chromatography before refolding into histone octamers and purifying by size exclusion chromatography. Nucleosomes were assembled by gradient dialysis against TE buffer at 4 °C overnight with 147 bp core Widom 601 DNA, with or without a a 60 bp-overhang (underlined), as indicated 5’- CTGGAGAATCCCGGTGCCGAGGCCGCTCAATTGGTCGTAGACAGCTCTAGCACCG CTTAAACGCACGTACGCGCTGTCCCCCGCGTTTTAACCGCCAAGGGGATTACTCC CTAGTCTCCAGGCACGTGTCAGATATATACATCCTGT<u>GCATGTATTGAACAGCGAC</u> <u>CTTGCCGGTGCCAGTCGGATAGTGTTCCGAGCTCCCACTCT</u>-3’ ^63^. For remodeling assays, the reaction buffer contained 20 mM Tris-HCl, pH 8.0, 75 mM NaCl, 2 mM MgCl2, and 1 mM ATP. After adding DDM1 to Widom 601 + 60 bp nucleosome samples in a 2:1 ratio, the reactions were incubated at 25 °C and stopped by addition of 5 mM EDTA and excess plasmid DNA. Reaction samples were resolved by 6% native PAGE (37.5:1 acrylamide:bis-acrylamide) run in 0.5× TBE buffer and stained with SYBR Gold (Thermo Fisher Scientific). For gel shift assays, nucleosomes were assembled with 147 bp core Widom 601 DNA. In brief, DDM1 was mixed with nucleosomes in the binding buffer (20 mM Tris-HCl, pH 8.0, 75 mM NaCl, 2 mM MgCl2) and incubated for 30 min. The DDM1- nucleosome complex samples were resolved by 6% native PAGE in 0.5 × TBE (29:1 acrylamide:bis-acrylamide).

Peptide binding assays of purified His-TEV-DDM1 or His-TEV-DDM1[133-end] were measured using a Monolith NT.115 Pico running MO Control version 1.6 (NanoTemper Technologies). Assays were performed in PBS-T (137 mM NaCl, 2.7 mM KCl, 10 mM Na2HPO4, 1.8m mM KH2PO4, 0.1% Tween-20) for DDM1 and PBS-T supplemented with 1mM ADP for DDM1[133-end]. His-label RED-tris-NTA (NanoTemper Technologies) labeled DDM1 or DDM1[133-end] (5 nM) was mixed with 16 serial dilutions of histone H4 peptides starting at 1 mM and loaded into microscale thermophoresis premium coated capillaries (NanoTemper Technologies). MST measurements were recorded at 23°C using 20% excitation power and 60% MST power. Measurements were performed in triplicate (except 133-end). Determination of the binding constant was performed using MO Affinity Analysis v.2.3.

#### Cryo-electron microscopy sample preparation

Purified DDM1 and reconstituted nucleosomes (H2A.W, H2B, H3.3, H4, and 147 bp DNA) were each desalted into binding buffer (10 mM HEPES, pH 7.5, 50 mM NaCl). DDM1 at 1.3 mg/mL and nucleosomes at ∼0.16 mg/mL were then mixed in a 4:1 molar ratio and incubated at room temperature for 10 minutes. DDM1-nucleosome complexes were cross-linked with 0.05% glutaraldehyde for 15 minutes then quenched by the addition of 2 mM Tris, pH 8.0. After five minutes at room temperature, the slowly-hydrolyzable ATP analog ATP-γ-S and MgCl2 were added to final concentrations of 1 mM and 2 mM, respectively. The reaction was incubated at 4°C overnight.

For cryo-EM grid preparation, 4 μL samples at approximately 0.35 mg/mL were applied to glow-discharged Quantifoil 0.6/1 300 μm mesh copper grids. After a 10 s incubation at 25 °C and 95% humidity, samples were blotted for 2.5 s then plunged into liquid ethane using an Automatic Plunge Freezer EM GP2 (Leica).

#### Cryo-electron microscopy data acquisition

Data were acquired on a Titan Krios transmission electron microscope (ThermoFisher) operating at 300 keV. EPU data collection software version 2.10.0.5 (ThermoFisher) was used to collect micrographs at a nominal magnification of 81,000x (1.1 Å/pixel) and defocus range of −1.0 to −2.2 μm. Dose-fractionated movies were collected using a K3 direct electron detector (Gatan) operating in electron counting mode. In total, 30 frames were collected over a 4.8 s exposure. The exposure rate was 14.8 e^-^/Å^2^/s, which resulted in a cumulative exposure of approximately 71.2 e^-^/Å^2^. In total, 8,165 micrographs were collected.

#### Cryo-electron microscopy data processing

Real-time image processing (motion correction, CTF estimation, and particle picking) was performed concurrently with data collection using WARP version 1.0.9 ^96^. Automated particle picking was performed with the BoxNet pretrained deep convolutional neural network bundle included with WARP that is implemented in TensorFlow. A particle diameter of 180 Å and a threshold score of 0.6 yielded 3,788,872 particle coordinates.

Of the particles collected during cryo-EM acquisition, nearly three-quarters were free nucleosomes. Classification and refinement were carried out in cryoSPARC v3.2.0+210831 ^97^. Initial 2D classification showed distinct classes of nucleosomes both bound to and independent of DDM1. To isolate the DDM1-bound nucleosome particles, 2D classes were first manually inspected. Classes that clearly showed the presence of DDM1—typically top views—were preferentially selected (497,127 particles) for *ab initio* reconstruction of four, 3D classes using a 200,000 particle subset. The resulting models (one of which showing DDM1-bound nucleosome), were then used for 3D heterogenous refinement with the full particle set. The resulting DDM1-bound nucleosome class was then taken for iterative rounds of homogenous refinement, non-uniform refinement, and further filtering using the refined reconstruction together with DDM1-free nucleosome decoy classes. The final non-uniform refined reconstruction was generated from 215,066 particles and had a resolution of 3.2 Å according to the gold standard FSC.

#### Molecular model building and refinement

An atomic model of the SWI/SNF nucleosome complex (PDB: 6UXW) ^118^ and the AlphaFold prediction of DDM1 ^119^ were used as initial references for model building in Coot version 0.9.2-pre ^120^. After the initial build was generated, density modification was performed using Resolve ^121^. Subsequent rounds of interactive model building and refinement were performed with Coot and Phenix version 1.19.2-4158-000 ^122^, respectively. Secondary structure restraints for both the protein (helix and β-strand) and DNA (base-stacking and base-pairing) were used throughout refinement. Structure validation was conducted by MolProbity version 4.5.1 ^123^. Data collection, processing, and model validation statistics are provided in Tables S7 and S8.

#### Molecular graphics

Figures of molecular models were generated using ChimeraX version 1.2.5 ^124^. Electrostatic surface calculations were performed by APBS ^100^ with a solvent ion concentration of 0.15 M at 298 K using the PARSE force field. Superpositioning of structural homologs was performed by the DALI server ^125^. Conservation analysis was performed using the Consurf server ^98^.

## Quantification and statistical analysis

The statistical details of analysis applied in this paper are provided alongside in the figure legends.

## Notes

### Competing Interest Statement

The authors have declared no competing interest.

### Summary of Updates

Changed license to CC BY

