## Supplementary Material for "Chromatin remodeling of histone H3 variants underlies epigenetic inheritance of DNA methylation"

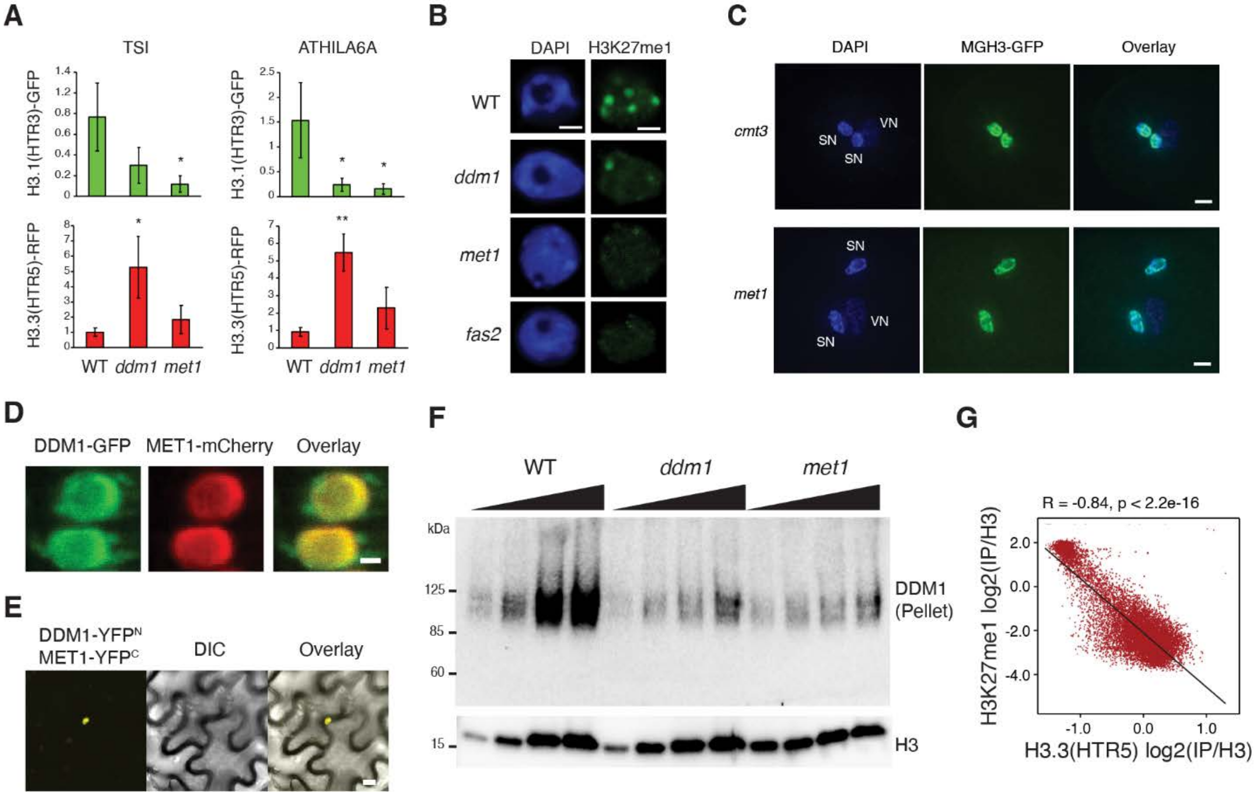


**Supplementary Figure S1. MET1 participates in histone remodeling by DDM1.** (A) ChIP-qPCR amplification of *TSI* (*ATHILA*) and *ATHILA6A* repeats in wild-type (WT), *ddm1*, and *met1* with 10-d-old seedling tissues. ChIP signals of H3.1(HTR3)-GFP and H3.3(HTR5)-RFP were normalized to H3. Error bars indicate standard deviations (biological replicates; n=3). P values of statistical difference with WT are shown above each mutant (one-way Anova adjusted with Tukey's Honest Significant Difference method; * p-value<0.05, ** p-value<0.01). (B) Immunofluorescence of H3.1-associated histone modification H3K27me1 in 3-week-old leaves of WT, *ddm1*, *met1*, and *fas2* (*caf-1*). DAPI was used for DNA staining. Scale bars indicate 2 µm. (C) Male Germline-specific Histone H3.3 MGH3-GFP localization in sperm nuclei of Arabidopsis pollen grains. DAPI staining was used to visualize vegetative (VN) and sperm nuclei (SN). Mislocalization to the nuclear periphery was observed in *met1* mutants, but not in *cmt3*. Scale bars indicate 2 µm. (D) Co-localization of DDM1-GFP and MET1-mCherry in the nucleus. Scale bar indicates 2 µm. (E) Bimolecular fluorescence complementation using DDM1 fused with N-terminal YFP (YFPN) and MET1 with C-terminal YFP (YFPC). Scale bar indicates 5 µm. Complementation is defined by the yellow nucleus. (F) Western blot analysis of endogenous DDM1 from the chromatin/pellet (P) fraction of WT, *ddm1*, and *met1* backgrounds. Anti-H3 was used as loading control. Serial dilutions (1:2) were made for each sample (gradient) indicating that both *ddm1* and *met1* mutants had between ½ and ¼ WT levels of chromatin-bound DDM1. (G) Genome-wide negative correlation between H3K27me1 (H3.1) and H3.3(HTR5)-RFP ChIP-seq in wild-type (Figure 1D). P and R values indicate statistical significance and Pearson correlation coefficients, respectively.

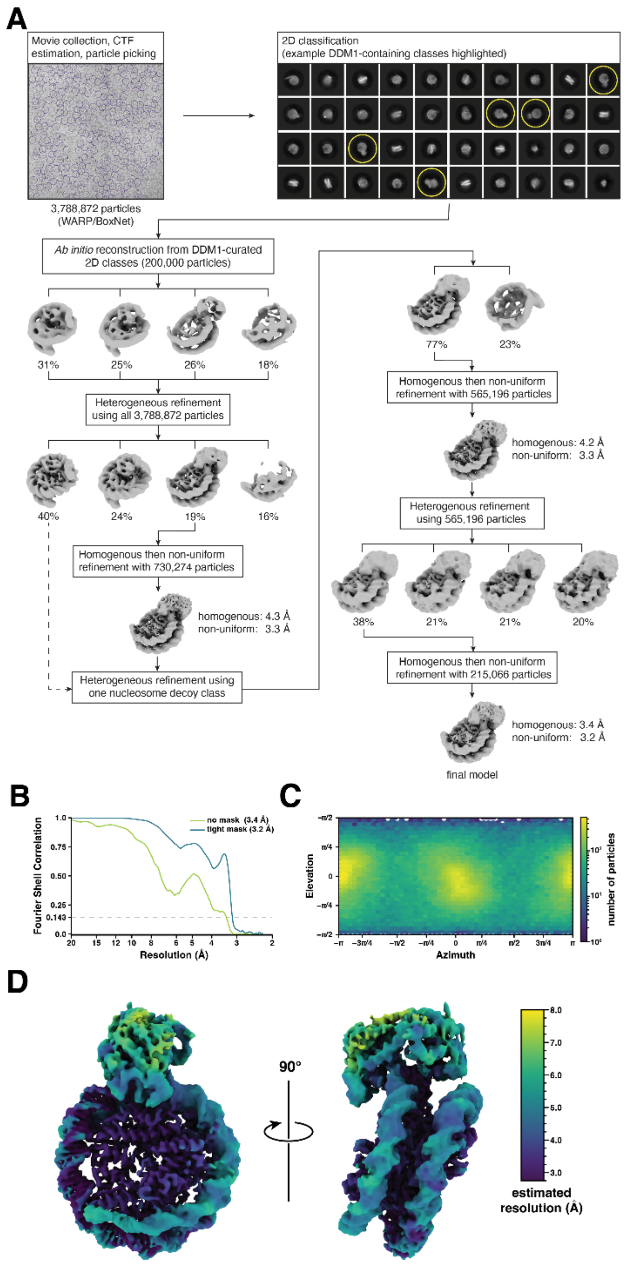


**Supplementary Figure S2. Cryo-EM data processing workflow and reconstruction metrics.** (A) Following cryo-EM movie collection, motion correction, averaging, CTF estimation, and particle picking were performed using WARP [96]. Example particle picks are shown as purple circles (top left). Particles were then imported into cryoSPARC [97] for 2D classification as well as 3D classification and refinement. Examples of DDM1-containing 2D classes, which were used to generate the ab initio models are highlighted with yellow circles (top right). Class distributions are indicated for each heterogenous refinement step. Reconstruction resolutions after homogenous and non-uniform refinement are indicated next to the corresponding models. (B) Fourier Shell Correlation (FSC) plots of the DDM1-nucleosome reconstruction using no mask (green) and a tight mask (blue). Resolution values at FSC 0.143 are indicated. (C) Angular distribution plot of reconstruction projections. The heat map indicates the number of particles per viewing angle. (D) The DDM1-nucleosome complex reconstruction, colored by estimated local resolution from cryoSPARC.

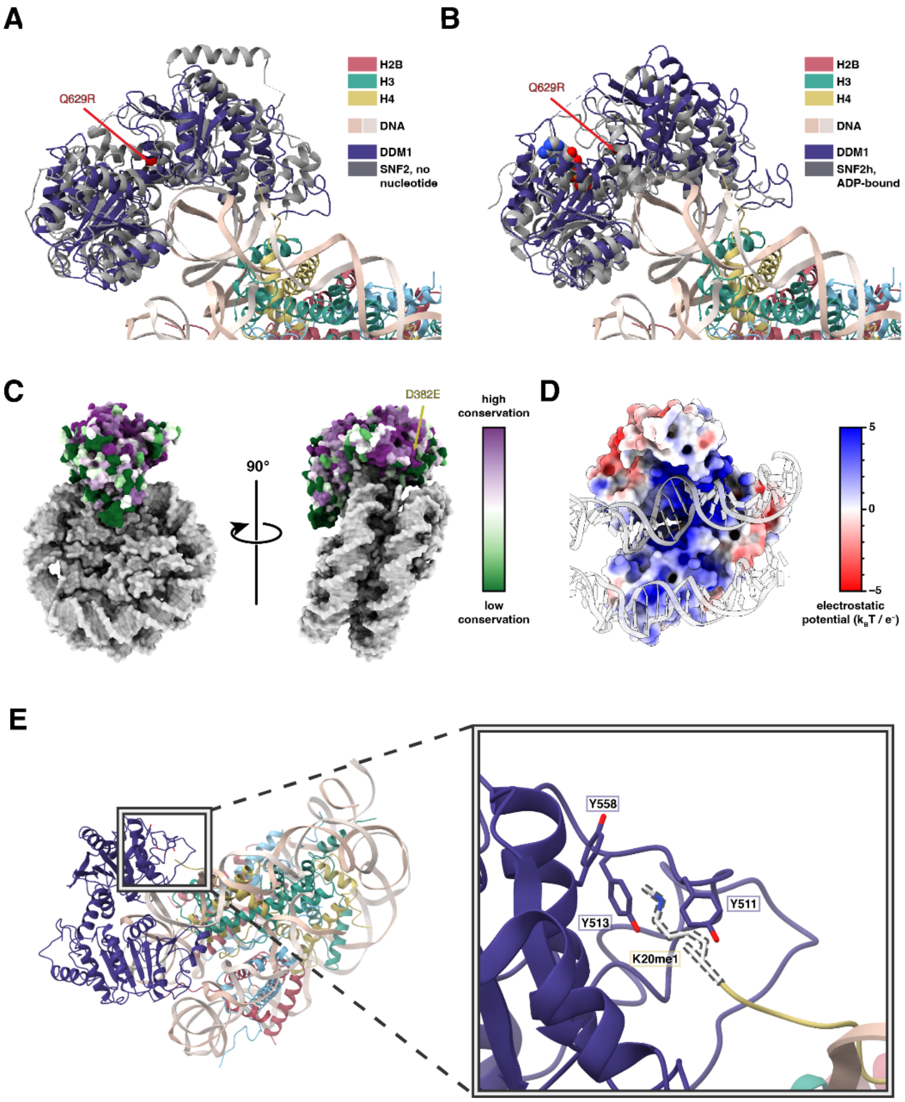


**Supplementary Figure S3. Structural comparison of DDM1 with Snf2 family remodelers.** The structures of (A) Snf2-bound nucleosomes in the absence of nucleotide (PDB code 5X0Y) and (B) Snf2h in the presence of ADP (PDB code 6NE3) (red) superimposed on the structure of DDM1 bound to nucleosome. The Q629R mutation in DDM1 is shown with a red arrow. Alignment was performed using only the nucleosome core particle for each structure. In the presence of bound ADP, the two domains appear in a more closed conformation than the nucleotide free state. The DDM1/nucleosome complex that was reconstructed represents the nucleotide free state. Note that the sample used for the Snf2h structure was prepared with ADP-$BeF_3$ but only ADP was observed in the density. (C) Surface representation of the DDM1-bound nucleosome colored according to degree of DDM1 conservation. Conservation scores were calculated using the Consurf server [98] among twenty manually-curated and highly related sequences—such as LSH and HELLS—aligned using Clustal Omega [99]. The D382A mutation in DDM1 is indicated with a yellow arrow. (D) The electrostatic potential of DDM1 (colored surface) displays a positively-charged groove along the DNA (grey cartoon) interface. Electrostatic surface calculations were performed by APBS [100] with a solvent ion concentration of 0.15 M at 298 K using the PARSE force field. (E) The tail of histone H4 extends toward DDM1 such that the residue K20 would be within striking distance of three aromatic residues in DDM1 forming an aromatic cage. The inset indicates a modeled mono-methylated lysine residue with a dashed outline.

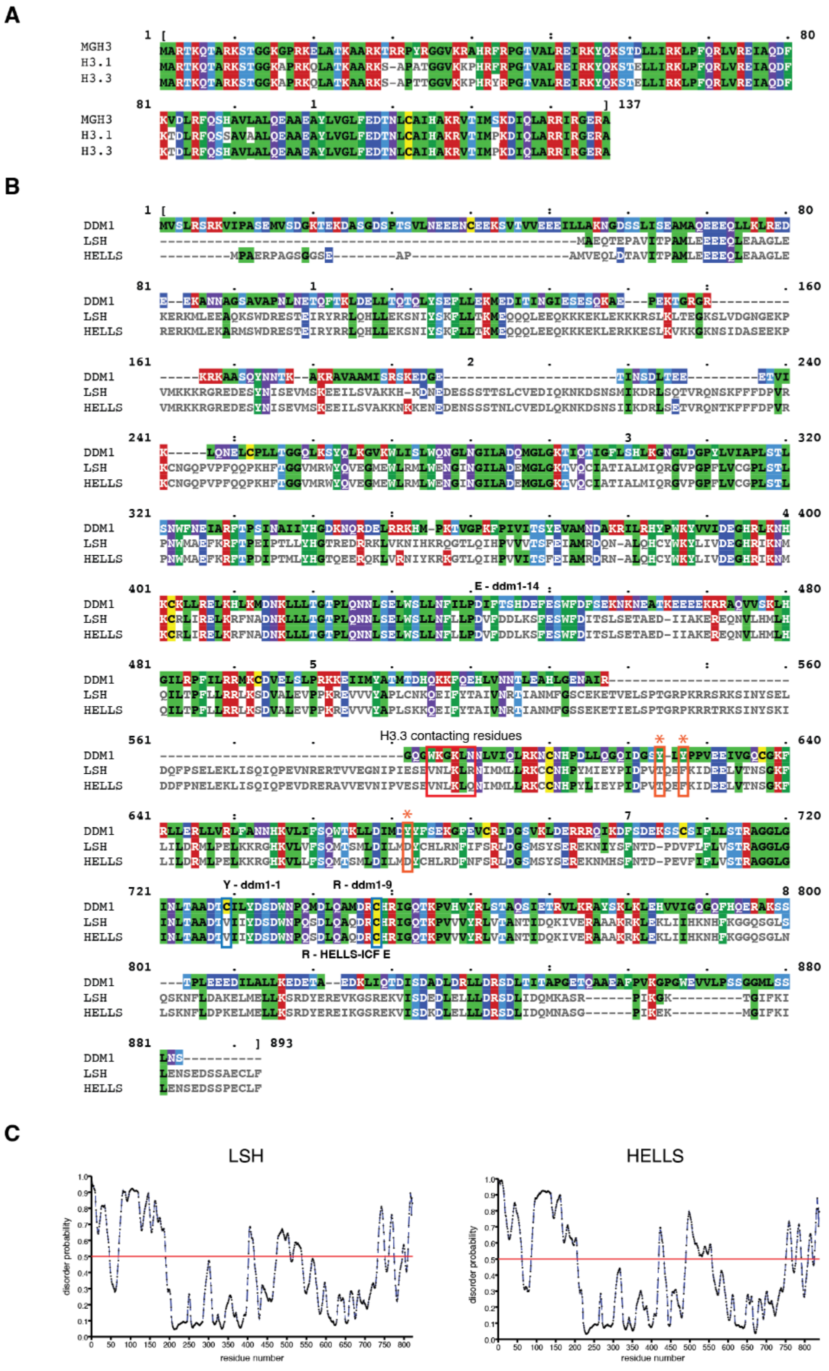


**Supplementary Figure S4. Amino acid sequence alignments.** (A) The sequence alignment of histone MGH3, H3.1, and H3.3 generated with MView [101]. (B) The sequence alignment of DDM1, LSH, and HELLS. H3.3 contacting residues (WKGKLN) of DDM1 are indicated with a red bar. Tyrosine residues Y511, Y513, and Y558 (DDM1 aromatic cage) are indicated with orange asterisks. Cysteine residues C615 and C634 (DDM1 disulfide bond) are indicated with blue asterisks. Three DDM1 hypomethylation mutations (*ddm1-1*, *ddm1-9* and *ddm1-14*) and one HELLS mutation (ICF proband family E) are indicated by substituted residues above and below the mutated location, respectively. Compared to DDM1, LSH has 90.6% coverage with 34.8% identity while HELLS has 92.8% coverage with 33.8% identity. (C) Intrinsically disordered regions of LSH and HELLS using PrDOS [102].

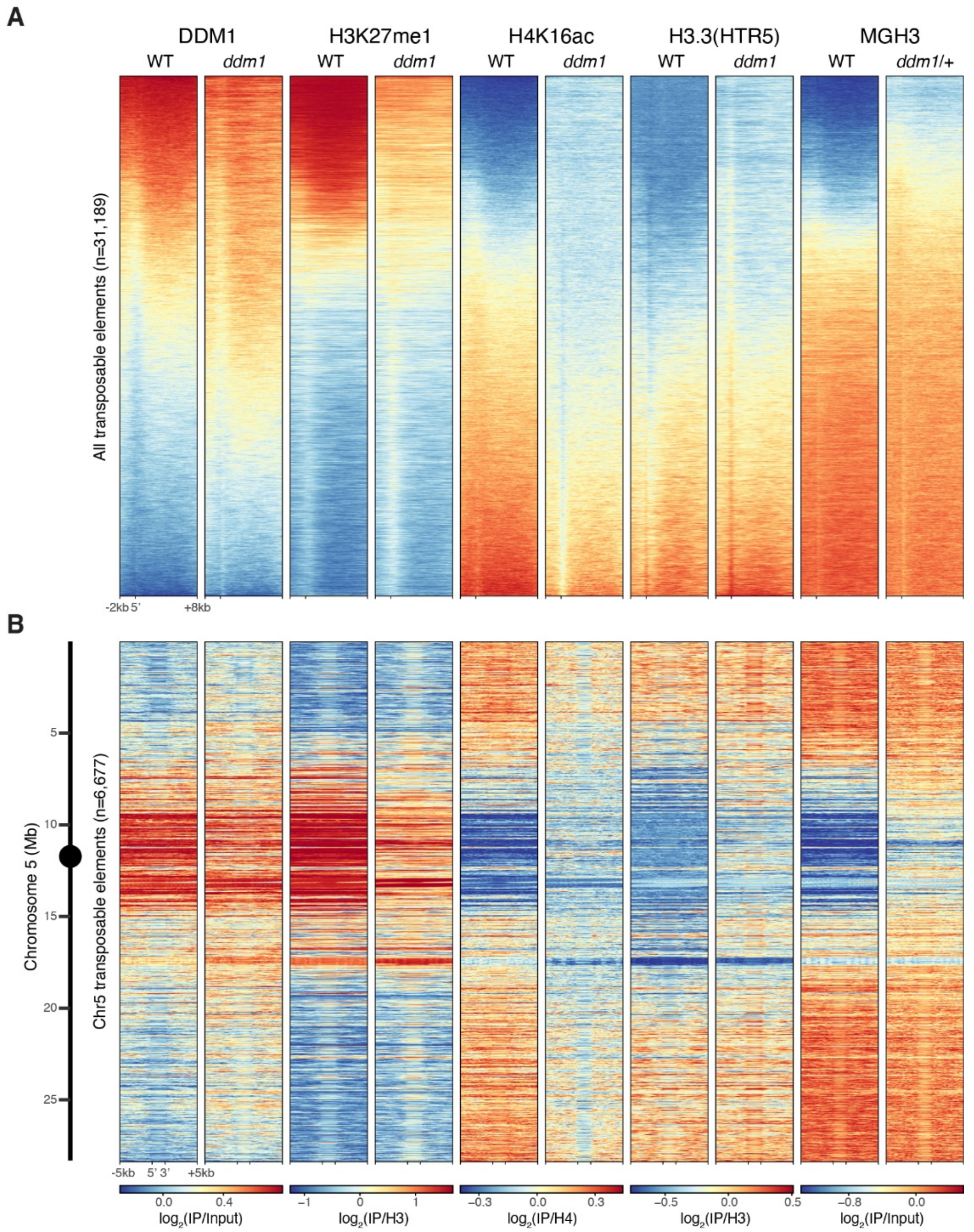


**Supplementary Figure S5. ChIP-seq data for all transposable elements in WT and *ddm1*.** (A) Heatmaps of DDM1, H3K27me1, H4K16ac, H3.3(HTR5) ChIP-seq of wild-type (WT) and *ddm1* genotypes, as well as MGH3 in pollen from WT and *ddm1*/+ plants, for all transposable elements annotated in TAIR10. Heatmaps were generated using Deeptools [95], where all 31,189 TEs were aligned by their 5' end with 2kb upstream and 8kb downstream with a binsize of 10bp, and sorted based on DDM1 levels in WT. (B) Similar heatmaps were generated using Deeptools, where the 6,677 TEs located on chromosome 5 were scaled to 2kb, represented with 5kb upstream and 5kb downstream with a binsize of 10bp. TEs were kept in order of their location on the chromosome, shown by the scale on the left hand-side. This view highlights that DDM1 preferentially targets peri-centromeric TEs in WT. Both heatmaps highlight correlation between DDM1 and H3K27me1 and anti-correlation with H4K16ac, H3.3 and MGH3 levels in both genotypes, as well as the loss of DDM1 and H3K27me1 from peri-centromeric TEs in *ddm1* accompanied by an increase in H4K16ac, H3.3 and MGH3.

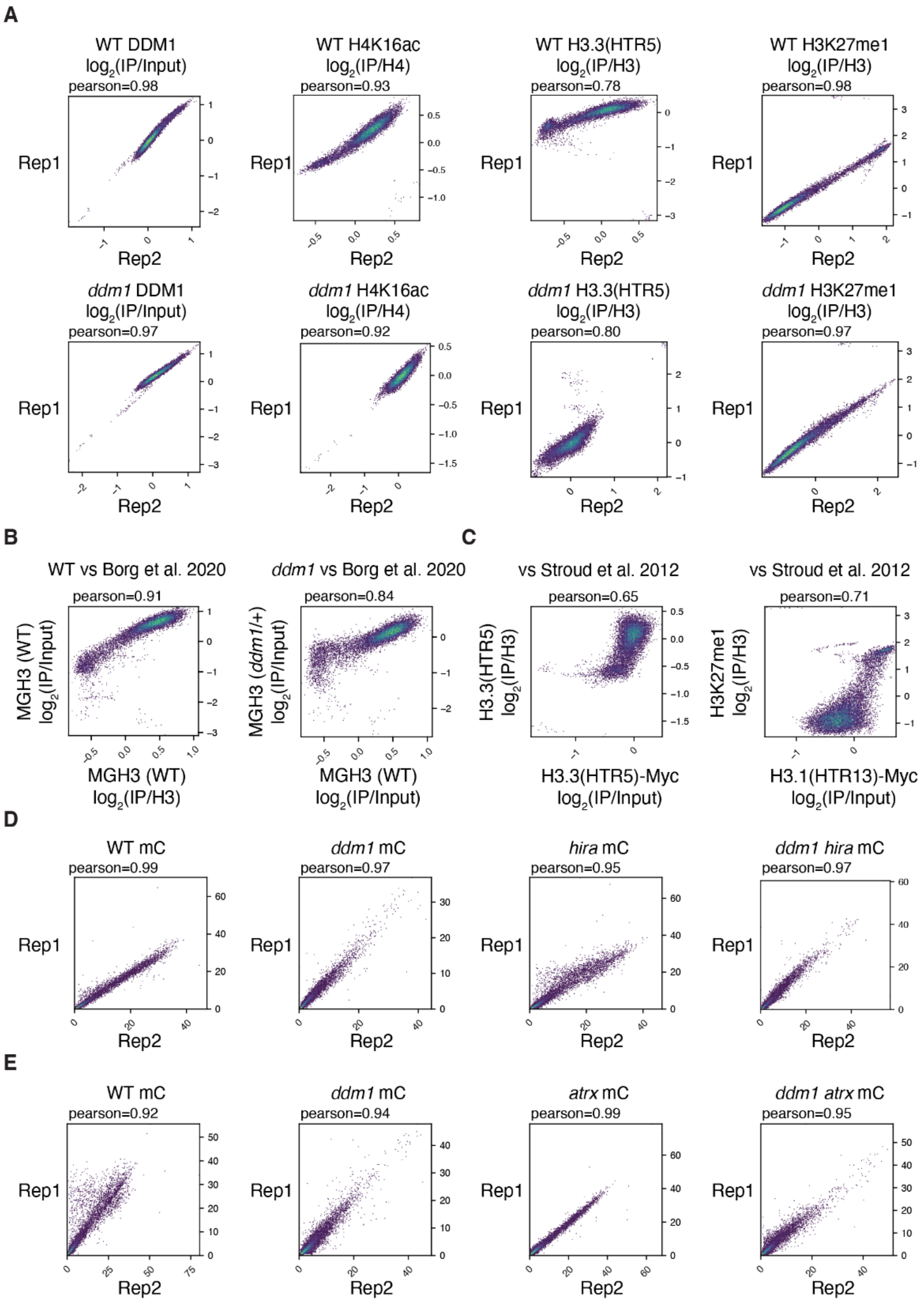


**Supplementary Figure S6. Correlations between ChIP-seq replicates and between WGBS replicates.** (A) Comparisons of DDM1, H4K16ac, H3.3(HTR5)-RFP and H3K27me1 ChIP-seq data between replicates of each genotype. Pearson correlations are shown. (B) Comparisons of MGH3 in WT and ddm1/+ pollen with previously published MGH3 ChIP-seq [47]. (C) Comparisons of H3.3(HTR5)-RFP and H3K27me1 ChIP-seq in WT with previously published H3.3(HTR5)-Myc and H3.1(HTR13)-Myc, respectively [40]. (A-C) Each replicated IP has been normalized to its respective input. (D, E) Comparisons of DNA methylation levels in each replicate for all genotypes grown and processed at the same time for group A (D) and group B (E), respectively.

**Supplementary Video S1 -** Live imaging of DDM1-GFP and H3.3(HTR5)-RFP during transition from M phase to G1 phase.

**Supplementary Video S2 -** Live imaging of DDM1-GFP and H3.3(HTR5)-RFP during transition from G2 phase to M phase.

**Supplementary Table S1 -** Coordinates of hypermethylated differentially methylated regions in ddm1 hira vs ddm1, Related to Figure 2 (.xlsx).

**Supplementary Table S2 -** Coordinates of stable and revertant DMRs in ddm1 identified in Colomé-Tatché et al., 2012, and random control regions, Related to Figure 6 (.xlsx).

**Supplementary Table S3. List of mutants in this study, Related to STAR Methods**

| Mutants | Mution type | Notes |
|---|---|---|
| *ddm1-2* | EMS mutant (nucleotide: G to A) | Hypomorphic; the splicing defect leads to a deletion, a frameshift and premature translation termination; DNA hypomethylation |
| *ddm1-10* | T-DNA (SALK_093009) | T-DNA insertion into exon region; used for crossing with MGH3-GFP |
| *met1-1* | EMS mutant (amino acid: P 1300 S) | Hypomorphic; amino acid substitution in catalytic domain; DNA hypomethylation; delayed flowering |
| *met1-7* | T-DNA (SALK_076522) | T-DNA insertion into exon region; used for crossing with MGH3-GFP |
| *cmt3-11* | EMS mutant | Single nucleotide substitution leading to a nonsense mutation; used for crossing with MGH3-GFP |
| *fas2-4* | T-DNA (SALK_033228) | Null; no transcript detectable; fasciation |
| *hira-1* | WiscDsLox362H05 | Reduced fertility |
| *atrx-2* | SAIL_861_B04 | Reduced fertility |

### Supplementary Table S4. Primer sequences, Related to STAR Methods

#### Cloning primers

| Name | Primer sequence (5' to 3') | Locus | Note |
|---|---|---|---|
| DDM1_attB1 | GGGGACAAGTTTGTACAAAAAAGCAGGCT ATTACTAATTGTGTCGACAAATCC | AT5G66750 | for cloning into pDONR221 |
| DDM1_attB2 | GGGGACCACTTTGTACAAGAAAGCTGGGT AATCCCAAATCCAAAACATAAGATC | AT5G66750 | for cloning into pDONR221 |
| MET1_attB1 | GGGGACAAGTTTGTACAAAAAAGCAGGCT TCATGGTAAAATGTTAGTTCTCG | AT5G49160 | for cloning into pDONR221 |
| MET1_attB2 | GGGGACCACTTTGTACAAGAAAGCTGGGT CTGGACAAACTTTATTTCGAC | AT5G49160 | for cloning into pDONR221 |

#### Genotyping

| Name | Primer sequence (5' to 3') | Locus | Note |
|---|---|---|---|
| ddm1-2 CAPS F | GTTGGACAGTGTGGTAAATTCCGCT | AT5G66750 | RsaI digestion |
| ddm1-2 CAPS R | GAGCTACGAGCCATGGGTTTGTGAAACGTA | AT5G66750 | RsaI digestion |
| ddm1-10 F | GCAAGCCATGGACAGATGCCACAG | AT5G66750 | |
| ddm1-10 R | CAGAGGGCCAATTGTTTTCATCAC | AT5G66750 | |
| met1-1 F | CTCTTTAGTAGAAGTTGGCATG | AT5G49160 | HaeIII digestion |
| met1-1 R | ATATGTATGTATAGATATTTTCTCC | AT5G49160 | HaeIII digestion |
| hira-1 LB | CTACTAAAATTTGAGGCCGGG | AT3G44530 | |
| hira-1 RB | GAGAGTCACTGTTTTGGCTGG | AT3G44530 | |
| atrx-2 LB | AGGAACCCTCACAGCTTCTTC | AT1G08600 | |
| atrx-2 RB | TCACATGGATGGCTTCTTTTC | AT1G08600 | |
| HTR3-GFP-F | TAGTGCAGTCGCAGCTCTTC | AT3G27360 | |
| HTR3-GFP-R | TTTGTACAAGAAAGCTGGGTCAG | AT3G27360 | |

#### ChIP-qPCR

| Name | Primer sequence (5' to 3') | Locus | Note |
|---|---|---|---|
| TSI F | ATCCAGTCCGAAGAACGCGAACTA | | |
| TSI R | TCACTTGTGAGTGTTCGTGAGGTC | | |
| ATHILA6A F | ACAGGAAGTGGGCGCACACC | AT5G32511 | |
| ATHILA6A R | CTCACAACGACGCAAGTGATCT | AT5G32511 | |

#### ATPase assay substrates

| Name | Primer sequence (5' to 3') | Locus | Note |
|---|---|---|---|
| Widom601 0N60 F | AGAGTGGGAGCTCGGAACACTATCCGAC | | |
| Widom601 0N60 R | CTGGAGAATCCCGGTGCC | | |

### Supplementary Table S5. ChIP-sequencing libraries metrics, Related to STAR Methods

| Genotype | Sample | IP or Control | Replicate | Total reads | All mapped reads | (% total) | Uniquely mapped reads | (% total) |
|---|---|---|---|---|---|---|---|---|
| WT | DDM1 | IP | Rep1 | 40,515,724 | 37,212,763 | 91.85% | 21,468,053 | 52.99% |
| WT | DDM1 | Input | Rep1 | 47,290,771 | 45,973,121 | 97.21% | 32,105,406 | 67.89% |
| WT | DDM1 | IP | Rep2 | 34,645,829 | 32,569,880 | 94.01% | 19,276,714 | 55.64% |
| WT | DDM1 | Input | Rep2 | 51,279,688 | 49,817,225 | 97.15% | 34,561,985 | 67.40% |
| *ddm1* | DDM1 | IP | Rep1 | 50,102,086 | 47,183,497 | 94.17% | 27,125,034 | 54.14% |
| *ddm1* | DDM1 | Input | Rep1 | 39,982,483 | 39,028,473 | 97.61% | 26,984,304 | 67.49% |
| *ddm1* | DDM1 | IP | Rep2 | 49,577,213 | 43,872,422 | 88.49% | 24,804,809 | 50.03% |
| *ddm1* | DDM1 | Input | Rep2 | 33,380,544 | 32,612,138 | 97.70% | 22,752,619 | 68.16% |
| WT | H3K27me1 | IP | Rep1 | 30,078,663 | 29,656,796 | 98.60% | 11,522,922 | 38.31% |
| WT | H3K27me1 | H3 | Rep1 | 22,622,734 | 22,483,854 | 99.39% | 16,142,362 | 71.35% |
| WT | H3K27me1 | IP | Rep2 | 50,994,633 | 40,503,003 | 79.43% | 13,950,016 | 27.36% |
| WT | H3K27me1 | H3 | Rep2 | 60,442,954 | 59,862,967 | 99.04% | 45,680,564 | 75.58% |
| *ddm1* | H3K27me1 | IP | Rep1 | 26,031,998 | 25,885,357 | 99.44% | 18,435,013 | 70.81% |
| *ddm1* | H3K27me1 | H3 | Rep1 | 28,806,975 | 28,258,079 | 98.09% | 10,926,824 | 37.93% |
| *ddm1* | H3K27me1 | IP | Rep2 | 37,494,713 | 28,836,811 | 76.91% | 9,554,115 | 25.48% |
| *ddm1* | H3K27me1 | H3 | Rep2 | 52,356,409 | 51,800,730 | 98.94% | 38,115,613 | 72.80% |
| WT | H4K16ac | IP | Rep1 | 29,867,512 | 29,113,254 | 97.47% | 19,477,351 | 65.21% |
| WT | H4K16ac | H4 | Rep1 | 32,025,461 | 31,151,067 | 97.27% | 17,149,656 | 53.55% |
| WT | H4K16ac | IP | Rep2 | 49,958,674 | 48,846,091 | 97.77% | 33,687,425 | 67.43% |
| WT | H4K16ac | H4 | Rep2 | 49,482,528 | 48,239,575 | 97.49% | 26,621,503 | 53.80% |
| *ddm1* | H4K16ac | IP | Rep1 | 44,578,514 | 43,566,178 | 97.73% | 28,553,612 | 64.05% |
| *ddm1* | H4K16ac | H4 | Rep1 | 59,698,668 | 20,431,979 | 34.23% | 12,114,825 | 20.29% |
| *ddm1* | H4K16ac | IP | Rep2 | 40,697,481 | 39,804,154 | 97.81% | 26,494,041 | 65.10% |
| *ddm1* | H4K16ac | H4 | Rep2 | 33,698,770 | 32,821,242 | 97.40% | 17,607,160 | 52.25% |
| WT | HTR5 | IP | Rep1 | 89,816,986 | 89,262,349 | 99.38% | 74,274,600 | 82.70% |
| WT | HTR5 | H3 | Rep1 | 43,362,049 | 42,608,947 | 98.26% | 33,866,720 | 78.10% |
| WT | HTR5 | IP | Rep2 | 47,569,522 | 47,259,571 | 99.35% | 40,618,380 | 85.39% |
| WT | HTR5 | H3 | Rep2 | 62,187,338 | 61,578,996 | 99.02% | 46,846,643 | 75.33% |
| *ddm1* | HTR5 | IP | Rep1 | 91,478,171 | 90,971,481 | 99.45% | 68,533,360 | 74.92% |
| *ddm1* | HTR5 | H3 | Rep1 | 43,901,988 | 43,551,892 | 99.20% | 31,424,017 | 71.58% |
| *ddm1* | HTR5 | IP | Rep2 | 88,251,305 | 87,728,045 | 99.41% | 67,051,725 | 75.98% |
| *ddm1* | HTR5 | H3 | Rep2 | 77,052,248 | 76,533,910 | 99.33% | 51,185,737 | 66.43% |
| WT | MGH3 | IP | Rep1 | 114,691,469 | 110,674,811 | 96.50% | 97,134,182 | 84.69% |
| WT | MGH3 | Input | Rep1 | 42,239,090 | 39,899,471 | 94.46% | 25,189,691 | 59.64% |
| *ddm1* /+ | MGH3 | IP | Rep1 | 28,958,163 | 23,043,589 | 79.58% | 17,882,624 | 61.75% |
| *ddm1* /+ | MGH3 | Input | Rep1 | 25,645,477 | 24,623,469 | 96.01% | 14,422,150 | 56.24% |

### Supplementary Table S6. Bisulfite sequencing libraries metrics, Related to STAR Methods

| Genotype | Group | Replicate | Total raw reads | All mapped reads | (% total) | unique alignments | (% total) | deduplicated | (% unique) | Cytosine covered | Average coverage | Non conversion rate (% mC/C in Pt) |
|---|---|---|---|---|---|---|---|---|---|---|---|---|
| WT | A | Rep1 | 32,230,036 | 30,942,651 | 96.01% | 24,745,052 | 76.78% | 11,017,686 | 44.52% | 90.41% | 8.17 | 0.269861 |
| WT | A | Rep2 | 53,613,049 | 52,389,181 | 97.72% | 42,053,916 | 78.44% | 9,566,292 | 22.75% | 88.18% | 6.78 | 0.51281 |
| *hira* | A | Rep1 | 17,362,101 | 16,566,981 | 95.42% | 12,957,483 | 74.63% | 5,285,057 | 40.79% | 77.90% | 3.93 | 0.270725 |
| *hira* | A | Rep2 | 44,991,888 | 42,569,624 | 94.62% | 33,798,990 | 75.12% | 4,608,590 | 13.64% | 78.21% | 3.22 | 0.706732 |
| *ddm1hira* | A | Rep1 | 21,758,809 | 17,461,284 | 80.25% | 13,837,031 | 63.59% | 5,419,437 | 39.17% | 77.38% | 3.94 | 0.287453 |
| *ddm1hira* | A | Rep2 | 48,425,900 | 36,805,003 | 76.00% | 28,725,572 | 59.32% | 8,623,617 | 30.02% | 84.11% | 6.16 | 0.548739 |
| *ddm1* | A | Rep1 | 27,502,435 | 26,294,284 | 95.61% | 20,404,561 | 74.19% | 7,057,712 | 34.59% | 83.48% | 5.30 | 0.266461 |
| *ddm1* | A | Rep2 | 37,878,589 | 37,470,061 | 98.92% | 29,204,434 | 77.10% | 9,163,719 | 31.38% | 83.05% | 6.49 | 0.537523 |
| WT | B | Rep1 | 30,561,004 | 24,364,737 | 79.72% | 18,151,083 | 59.39% | 9,660,873 | 53.22% | 77.57% | 7.07 | 0.589116 |
| WT | B | Rep2 | 34,550,917 | 30,417,035 | 88.04% | 22,452,116 | 64.98% | 13,366,408 | 59.53% | 79.20% | 9.83 | 0.587723 |
| *ddm1atrx* | B | Rep1 | 23,525,857 | 19,197,760 | 81.60% | 14,136,528 | 60.09% | 8,880,690 | 62.82% | 71.88% | 6.43 | 0.595414 |
| *ddm1atrx* | B | Rep2 | 37,170,647 | 35,040,723 | 94.27% | 24,906,077 | 67.00% | 14,078,236 | 56.53% | 79.15% | 10.33 | 0.667431 |
| *ddm1* | B | Rep1 | 36,520,861 | 35,544,986 | 97.33% | 26,379,181 | 72.23% | 10,534,593 | 39.94% | 78.73% | 7.54 | 0.621665 |
| *ddm1* | B | Rep2 | 20,358,084 | 19,257,875 | 94.60% | 14,528,944 | 71.37% | 6,440,673 | 44.33% | 73.49% | 4.70 | 0.923219 |
| *atrx* | B | Rep1 | 39,661,456 | 39,246,987 | 98.95% | 29,154,984 | 73.51% | 7,359,080 | 25.24% | 79.63% | 5.21 | 0.845968 |
| *atrx* | B | Rep2 | 38,694,008 | 38,125,949 | 98.53% | 28,077,230 | 72.56% | 9,110,947 | 32.45% | 79.89% | 6.57 | 0.661121 |

**Supplementary Table S7. Cryo-EM data collection and reconstruction statistics for the DDM1-nucleosome complex, Related to STAR Methods**

| | |
|---|---|
| **Data collection** | |
| *Microscope* | Titan Krios |
| *Voltage (keV)* | 300 |
| *Magnification* | 81,000× |
| *Defocus range (μm)* | −1.0 to −2.2 |
| *Detector* | K3 |
| *Pixel Size (Å)* | 1.1 |
| *Total exposure ($e^-/Å^2$)* | 71.2 |
| *Exposure rate ($e^-/Å^2$/sec)* | 14.8 |
| *Exposure per frame ($e^-/Å^2$)* | 2.37 |
| *Micrographs collected* | 8,165 |

| | |
|---|---|
| **Initial processing** | |
| *Micrographs used* | 7,811 |
| *Initial particles* | 3,788,872 |

| | |
|---|---|
| **Reconstruction** | |
| *Final particles* | 215,066 |
| *Symmetry* | C1 |
| *Map sharpening B-factor ($Å^2$)* | 57.8 |
| *Half maps resolution (unmasked / masked)* | |
| FSC 0.143 | 3.4 / 3.2 |

**Supplementary Table S8. Model refinement and validation statistics for the DDM1-nucleosome complex, Related to STAR Methods**

Statistics are provided for the full model as well as the individual octamer, DNA and DDM1 components.

| **Refinement** | **Full model** | **Octamer** | **DNA** | **DDM1** |
|---|---|---|---|---|
| *Protein residues* | 1217 | 749 | —— | 468 |
| *Nucleic acid residues* | 282 | —— | 282 | —— |
| *Model resolution (unmasked / masked)* | | | | |
| FSC 0.5 | 3.2 / 3.2 | | | |
| FSC 0.143 | 2.8 / 2.8 | | | |
| *Map correlation coefficients* | | | | |
| CC mask | 0.74 | 0.78 | 0.73 | 0.6 |
| CC box | 0.65 | —— | —— | —— |
| CC peaks | 0.65 | —— | —— | —— |
| CC volume | 0.72 | 0.73 | 0.72 | 0.58 |

| **Model geometry** | **Full model** | **Octamer** | **DNA** | **DDM1** |
|---|---|---|---|---|
| *Ramachandran plot* | | | | |
| Outliers (%) | 0 | 0 | —— | 0 |
| Allowed (%) | 0.67 | 0 | —— | 1.72 |
| Favored (%) | 99.33 | 100 | —— | 98.28 |
| *Ramachandran plot Z-score* | | | | |
| Whole | −0.32 ± 0.23 (N = 1197) | 0.18 ± 0.30 (N = 733) | —— | −1.11 ± 0.36 (N = 464) |
| Helix | 0.25 ± 0.19 (N = 681) | 0.72 ± 0.23 (N = 492) | —— | -0.96 ± 0.34 (N = 189) |
| Sheet | 1.72 ± 0.73 (N = 55) | —— | —— | 1.72 ± 0.73 (N = 55) |
| Loop | −1.05 ± 0.26 (N = 461) | −1.00 ± 0.36 (N = 241) | —— | −1.10 ± 0.38 (N = 220) |
| *CaBLAM outliers (%)* | 0.4 | 0.3 | —— | 0.7 |
| *Rotamers* | | | | |
| Poor (%) | 0.1 | 0 | —— | 0.24 |
| Favored (%) | 98.28 | 98.72 | —— | 97.62 |
| *R.M.S. deviations* | | | | |
| Bond lengths (Å) | 0.004 | 0.004 | 0.004 | 0.004 |
| Bond angles (°) | 0.759 | 0.757 | 0.716 | 0.837 |
| *Geometry outliers* | | | | |
| $C_\alpha$ deviations (%) | 0.08 | 0 | —— | 0.22 |
| $C_\beta$ deviations | 0 | 0 | —— | 0 |
| Bad bonds | 0 / 9,894 | 0 / 5,991 | 0 / 6,484 | 0 / 3,903 |
| Bad angles | 0 / 13,296 | 0 / 8,040 | 0 / 10,003 | 0 / 5,256 |
| Cis prolines | 0 / 39 | 0 / 24 | —— | 0 / 15 |
| Chiral volume outliers | 0 / 2633 | 0 / 931 | 0 / 1,128 | 0 / 574 |
| *Clashscore (all atoms)* | 3.22 | 1.97 | 2.79 | 5.57 |
| *MolProbity score* | 1.11 | 0.96 | —— | 1.3 |
| *B-factors ($Å^2$) (min / max / mean)* | | | | |
| Protein | 12.6 / 98.4 / 45.8 | 12.6 / 93.9 / 30.7 | —— | 36.1 / 98.4 / 69.2 |
| Nucleotide | 28.84 / 201.26 / 87.65 | —— | 28.8 / 201.3 / 87.7 | —— |
